# Adrenomedullin Restores Mitochondrial Bioenergetics and Rescues Interneuron Phenotypes in Human Models of 22q11.2 Deletion Syndrome

**DOI:** 10.64898/2026.05.24.726075

**Authors:** Dhriti Nagar, Jongbin Choi, Zunsong Hu, Li Li, Madison S. House, Seyeon Park, Bailey Abel, Anushka Nikhil, Meghali Aich, Saw Htun, Katja G. Weinacht, Zhaohui Gu, Anca M. Pasca

## Abstract

22q11.2 deletion syndrome (22q11.2DS), also known as DiGeorge syndrome, is the strongest genetic risk factor for schizophrenia, yet the cellular mechanisms underlying this vulnerability remain incompletely understood. Using iPSC-derived human subpallial organoids (hSOs) and forebrain assembloids (hFAs), we identified key 22q11.2DS cellular and molecular phenotypes in migrating cortical interneurons and rescued them with Adrenomedullin (ADM), a 52-amino-acid peptide hormone that acts through the CLR/RAMP2 receptor complex. Specifically, we discover that 22q11.2DS interneurons exhibit impaired migration patterns, and that this is associated with excessive mitochondrial fragmentation, reduced oxidative phosphorylation, and dysregulated calcium signaling. Transcriptomic analysis reveals a glycolytic shift and widespread downregulation of oxidative phosphorylation genes. Next, we identify ADM as a potent rescue agent that restores mitochondrial morphology, membrane potential, respiration, calcium homeostasis, actin retrograde flow, and interneuron migration. Mechanistically, we find that ADM signals through two parallel pathways: PKA-mediated DRP1 Ser637 phosphorylation, which promotes mitochondrial fusion, and PLC-mediated calcium signaling, which normalizes actin dynamics. These phenotypes and their rescue are recapitulated in primary developing human cortical tissue carrying the 22q11.2 deletion. Overall, our study establishes a mitochondrial-cytoskeletal axis as a novel biological mechanism underlying the alterations observed in neuronal development in human preclinical models of 22q11.2DS and identifies possible therapeutic molecular targets for 22q11.2DS-associated neuropsychiatric pathology.

**HIGHLIGHTS:**

- Inhibitory neurons in 22q11.2DS organoids and assembloids show mitochondrial fragmentation and impaired migration
- ADM rescues mitochondrial morphology, respiration, and interneuron migration
- PKA and PLC pathways mediate separate arms of ADM’s rescue mechanism
- Phenotypes and ADM rescue are validated in human cortical tissue

## INTRODUCTION

The 22q11.2 deletion syndrome (22q11.2DS), also called DiGeorge syndrome or velocardiofacial syndrome, is the most common microdeletion disorder in humans, occurring in about 1 in 4,000 live births^1–3^. The syndrome results from a hemizygous deletion of approximately 45 genes at the 22q11.2 locus and presents with a highly diverse clinical phenotype, including cardiac malformations, immune deficiency, palatal abnormalities, and neuropsychiatric symptoms^1,2,4–7^. One of the most clinically important features of 22q11.2DS is its strong link to neuropsychiatric illness: individuals with the deletion face a 25 to 30-fold increased risk of developing schizophrenia, making 22q11.2DS the strongest known genetic risk factor for this disorder^6,8–11^. It also increases the risk for attention-deficit/hyperactivity disorder (ADHD), autism spectrum disorders, intellectual disability, and anxiety disorders^9,12,13^. Despite ongoing research, the cellular and molecular mechanisms linking hemizygous loss of genes at 22q11.2 to the highly increased risk for neuropsychiatric diseases remain poorly understood, and this, in turn, hinders the efforts to identify effective therapeutics. Until recently, mechanistic and therapeutic investigation in neuropsychiatry, 22q11.2DS included, has focused largely on excitatory neurons, the most abundant neuronal cell type in the cerebral cortex. However, robust emerging literature in the neuropsychiatry field of research has brought cortical interneurons to the forefront as a cell type of particular interest, while a separate and parallel line of work has begun to position mitochondria as the likely central hub for the diverse cellular and molecular phenotypes in neuropsychiatry.

These scientific trends are highly relevant to 22q11.2DS for two main reasons. First, a hallmark characteristic of the 22q11.2 locus is the presence of multiple genes that encode proteins with established roles in mitochondrial function, and preclinical in vivo and in vitro studies have consistently reported mitochondrial dysfunction in models of 22q11.2DS^2,6,9,14–18^. Second, disrupted interneuron number, subtype composition, or placement has been implicated in the pathophysiology of schizophrenia, epilepsy, and other neuropsychiatric disorders¹¹. Although lack of tissue availability has precluded direct postmortem studies in 22q11.2DS, preclinical and clinical data points consistently to interneuron dysfunction. For example, histology data from individuals with non-22q11.2DS schizophrenia showed reduced GABAergic neuron density[, rodent models of 22q11.2DS identified aberrant interneuron migration and cortical localization²¹,²² and an iPSC-derived 22q11.2DS cortical assembloids models found decreased transition of interneurons from the subpallial to the pallial organoids after fusion²³,

Given the increasing scientific emphasis and evidence for both interneuron defects and mitochondrial dysfunction as major contributors to the pathology of neuropsychiatric conditions, and the emerging, yet limited data about these topics in 22q11.2DS, in this study, we focused on the in-depth investigation of the mechanistic link between cortical interneuron migration defects and mitochondrial dysfunction.

To model interneuron migration in 22q11.2DS, we used human forebrain assembloids (hFAs) derived from the fusion of human subpallial organoids (hSOs) and human cortical organoids (hCOs); this robust and reproducible platform enables in-depth cellular and molecular study of tangential interneuron migration toward the cortex^19–22^. Separately, we validated the main phenotypes and rescue findings using primary developing human cortical tissue with or without the 22q11.2 deletion.

First, we re-analyze the public dataset published by Walsh et al^42^ for 22q11.2DS human forebrain assembloids and identify that (i) genes associated with neuropsychiatric diseases are preferentially differentially expressed in inhibitory neurons in 22q11.2DS, and that the (ii) mitochondrial dysfunction also looks enriched in inhibitory neuronal lineages in 22q11.2DS.

Next, we generate hFAs and show that inhibitory interneurons from 22q11.2DS exhibit impaired calcium signaling and neuronal hyperexcitability, a phenotype previously also reported in cortical excitatory neurons from hCOs derived from individuals with 22q11.2DS^12^. Moreover, we demonstrate a direct association between abnormal interneuron migration patterns (after their transition from the hSO into the hCO), abnormal actin dynamics, mitochondrial fragmentation, and decreased oxidative phosphorylation in 22q11.2DS. Lastly, we show the rescue potential of adrenomedullin (ADM), a 52-amino-acid peptide hormone that acts through the CLR/RAMP2 receptor complex. We focused our rescue and mechanistic studies on ADM for two main reasons: (i) ADM is being explored as a therapeutic agent in sepsis, cardiovascular diseases and inflammatory bowel diseases due to its roles as an anti-inflammatory peptide^23,24^, and (ii) our previous study focused on defects in interneuron migration upon exposure to hypoxia^20^ identified ADM as a strong neuro-protective, thus expanding its clinical and therapeutic relevance to hypoxic brain injuries.

Mechanistically, we show that ADM acts through two parallel pathways in 22q11.2DS: the PKA-mediated phosphorylation of DRP1 at Ser637 promotes mitochondrial fusion, while PLC-mediated calcium signaling restores cytoskeletal dynamics in 22q11.2 interneurons.

To check whether these findings remain valid in a separate, even more relevant human system, we use primary human developing cortical tissue carrying 22q11.2 deletion, along with age-matched control tissue, and validate the major phenotypes and their rescue by ADM.

Overall, in this study, we describe and characterize interneuron migration pattern defects in 22q11.2DS, highlight a mitochondrial-cytoskeletal axis as a convergent disease mechanism, and suggest ADM and its related molecular pathways as potential therapeutic targets.

## RESULTS

### Analysis of a Published 22q11.2DS Assembloid Model Dataset Reveals Convergent Mitochondrial Dysregulation and Neuropsychiatric Risk Enrichment in Inhibitory Neurons Lineage

To identify convergent molecular signatures of the 22q11.2 deletion in an independently generated human cellular model, we first analyzed a publicly available single-cell RNA-seq dataset from 22q11.2DS brain assembloids containing both excitatory and inhibitory neurons from Walsh et al. (2026; GSE250482). Cross-referencing differentially expressed genes (DEGs) with published neuropsychiatric disorder risk gene sets revealed substantial overlap preferentially in inhibitory neurons across schizophrenia (SCZ; 65 genes, p < 0.001, Fisher’s exact test), bipolar disorder (BD; 47 genes, p < 0.001, Fisher’s exact test), autism spectrum disorder (ASD; 27 genes, p = 0.000425, Fisher’s exact test), and major depressive disorder (MDD; 18 genes, p = 0.000018, Fisher’s exact test). Interestingly, excitatory neuron overlap was not significant for any disorder in this dataset and ADHD-associated genes showed minimal overlap in either cell type (inhibitory: 1 gene, p = 0.639, Fisher’s exact test) (Figure 1A). These analyses suggest that GABAergic lineages might harbor a disproportionate share of the 22q11.2DS-associated neuropsychiatric transcriptional vulnerability.

**Figure 1.**
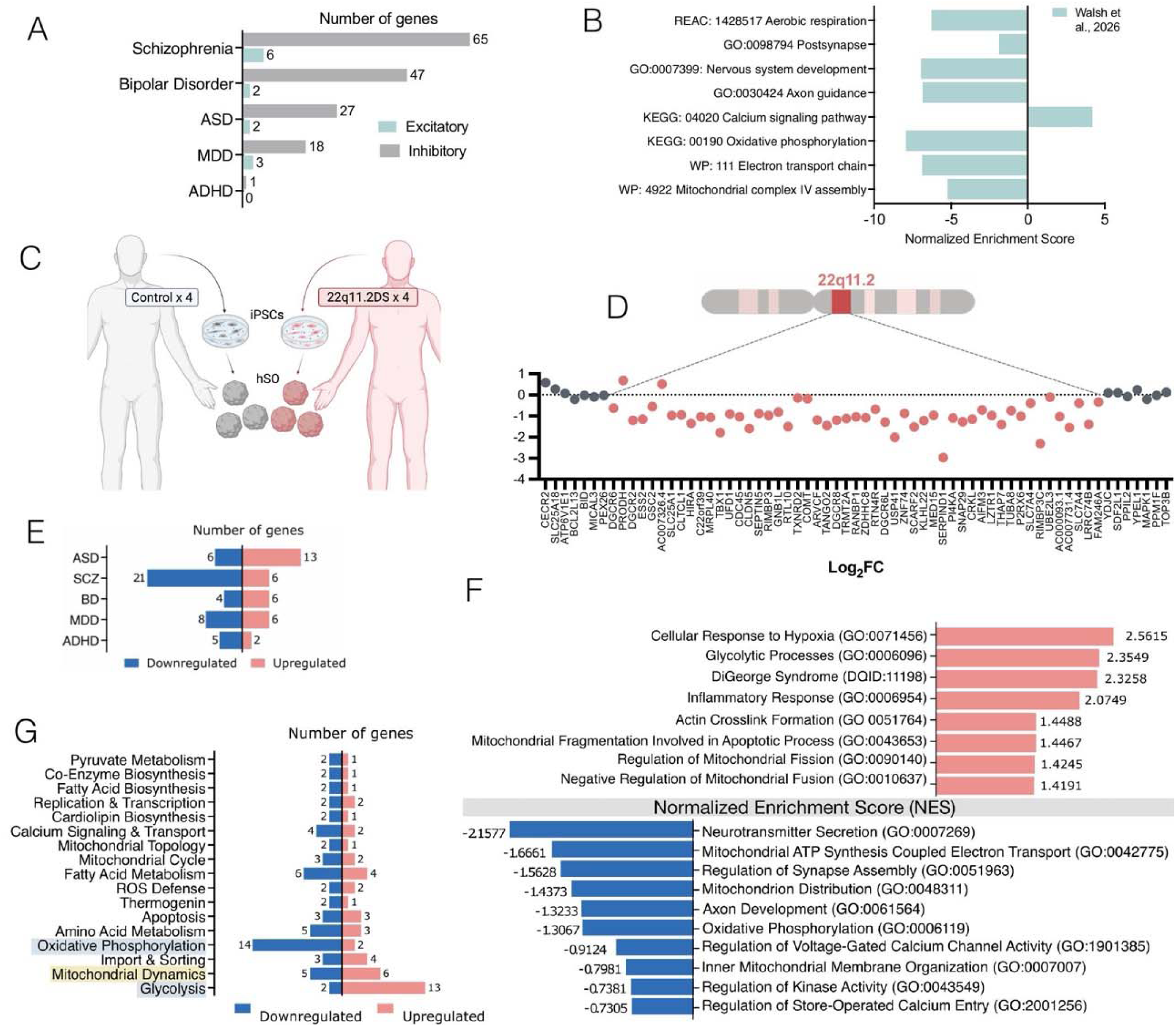
Vulnerability of inhibitory neurons in 22q11.2 deletion syndrome. Reanalysis of an independently published single-cell RNA-seq (scRNA-seq) datasets from 22q11.2DS brain assembloids reveals preferential vulnerability in inhibitory neurons for dysregulation of genes associated with neuropsychiatric conditions, mitochondrial dynamics and biogenesis genes. (A) Number of differentially expressed genes (DEGs) in inhibitory (grey) versus excitatory (teal) neurons from 22q11.2DS brain assembloids that overlap with neuropsychiatric disease-associated gene sets (Walsh et al., 2026). Inhibitory neurons show substantially greater overlap across all disorders examined: schizophrenia (SCZ; 65 genes, p < 0.001, Fisher’s exact test), bipolar disorder (BD; 47 genes, p < 0.001, Fisher’s exact test), autism spectrum disorder (ASD; 27 genes, p = 0.000425, Fisher’s exact test), and major depressive disorder (MDD; 18 genes, p = 0.000018, Fisher’s exact test). Excitatory neuron overlap was not significant for any disorder (SCZ: 6 genes, p = 0.219, Fisher’s exact test; BD: 2 genes, p = 0.586, Fisher’s exact test; ASD: 2 genes, p = 0.652, Fisher’s exact test; MDD: 3 genes, p = 0.070, Fisher’s exact test; ADHD: 0 genes, p = 1.0, Fisher’s exact test). ADHD-associated genes showed minimal overlap in either cell type (inhibitory: 1 gene, p = 0.639, Fisher’s exact test). One-sided Fisher’s exact test was performed using 600 inhibitory and 100 excitatory DEGs against a background of 22,116 genes. (B) Pathway enrichment analysis of 22q11.2DS (Walsh et al., 2026) DEGs. Enriched pathways include aerobic respiration (REAC: 1428517;-log₁₀pAdj = 2.177, Fisher’s exact test), postsynapse (GO: 0098794;-log₁₀pAdj = 10.502, Fisher’s exact test), nervous system development (GO: 0007399;-log₁₀pAdj = 3.754, Fisher’s exact test), axon guidance (GO: 0030424;-log₁₀pAdj = 2.578, Fisher’s exact test), calcium signaling pathway (KEGG: 04020;-log₁₀pAdj = 4.168, Fisher’s exact test), oxidative phosphorylation (KEGG: 00190;-log₁₀pAdj = 5.887, Fisher’s exact test), mitochondrial complex I assembly (WP: 4324, Fisher’s exact test), electron transport chain (WP: 111;-log₁₀pAdj = 6.879, Fisher’s exact test), and mitochondrial complex IV assembly (WP: 4922;-log₁₀pAdj = 5.225, Fisher’s exact test; the X-axis represents the normalized enrichment score (NES). (C) Schematic of experimental design. Four control and four 22q11.2DS patient-derived induced pluripotent stem cell (iPSC) lines were differentiated into hSOs. (D) Top: schematic of the 22q11.2 chromosomal locus. Bottom: log₂ fold change (Log₂FC) of individual genes across the 22q11.2 region in 22q11.2DS versus control hSOs from bulk RNA-seq. (E) Overlap of 22q11.2DS hSO DEGs with neuropsychiatric disorder risk gene sets: autism spectrum disorder (ASD), schizophrenia (SCZ), bipolar disorder (BD), major depressive disorder (MDD), and attention-deficit/hyperactivity disorder (ADHD). Blue bars indicate downregulated genes; red bars indicate upregulated genes. Numbers indicate overlapping DEG counts. Fisher’s exact test. (F) Top enriched Gene Ontology (GO) terms ranked by Normalized Enrichment Score (NES). Upregulated pathways (positive NES, red): Cellular Response to Hypoxia (GO:0071456, NES = +1.53, p = 0.008, GSEA pre-ranked), Glycolytic Process (GO:0006096, NES = +1.32, p < 0.001, GSEA pre-ranked), DiGeorge Syndrome (GO:0006952, NES = +1.24, p < 0.001, GSEA pre-ranked), Inflammatory Response (GO:0006954, NES = +0.97, p = 0.014, GSEA pre-ranked), Actin Crosslink Formation (GO:0051764, NES = +0.86, p = 0.030, GSEA pre-ranked), Mitochondrial Fragmentation Involved in Apoptotic Process (GO:0043653, NES = +0.60, p = 0.015, GSEA pre-ranked), Regulation of Mitochondrial Fission (GO:0090140, NES = +0.45, p = 0.019, GSEA pre-ranked), Negative Regulation of Mitochondrial Fusion (GO:0010637, NES = +0.44, p = 0.002, GSEA pre-ranked). Downregulated pathways (negative NES, blue): Neurotransmitter Complex (GO:0097299, NES = −2.08, p < 0.001, GSEA pre-ranked), Mitochondrial ATP Synthesis Coupled Electron Transport (GO:0042775, NES = −1.56, p < 0.001, GSEA pre-ranked), Regulation of Synapse Assembly (GO:0051963, NES = −1.52, p = 0.012, GSEA pre-ranked), Mitochondrion Distribution (GO:0048312, NES = −1.46, p = 0.0103, GSEA pre-ranked), Axon Development (GO:0061564, NES = −1.20, p = 0.0056, GSEA pre-ranked), Oxidative Phosphorylation (GO:0006119, NES = −1.08, p = 0.0069, GSEA pre-ranked), Regulation of Voltage-Gated Calcium Channel Activity (GO:1901385, NES = −0.93, p = 0.0037, GSEA pre-ranked), Inner Mitochondrial Membrane Organization (GO:0007007, NES = −0.80, p = 0.0241, GSEA pre-ranked), Regulation of Kinase Activity (GO:0043549, NES = −0.62, p = 0.041, GSEA pre-ranked), Regulation of Store-Operated Calcium Entry (GO:0002931, NES = −0.48, p = 0.009, GSEA pre-ranked). Nominal p-values from GSEA pre-ranked analysis. (G) Number of dysregulated genes per mitochondrial functional category in 22q11.2DS versus control hSOs. Categories include pyruvate metabolism, co-enzyme biosynthesis, fatty acid biosynthesis, replication and transcription, cardiolipin biosynthesis, calcium signaling and transport, mitochondrial topology, mitochondrial cycle, fatty acid metabolism, reactive oxygen species (ROS) defense, thermogenin, apoptosis, amino acid metabolism, oxidative phosphorylation, import and sorting, mitochondrial dynamics, and glycolysis. Red bars indicate upregulated genes; blue bars indicate downregulated genes. Glycolysis shows the most upregulated genes (13 up, 2 down); oxidative phosphorylation shows the most downregulated genes (14 down, 2 up). Further details on the DEGs are provided in Table S3. See also Figures S1 and S2.

Separately, pathway enrichment analysis of the same DEG sets revealed striking convergence on mitochondrial bioenergetics and neuronal connectivity (Figure 1B). The dataset showed pronounced negative enrichment of aerobic respiration (REAC:1428517), oxidative phosphorylation (KEGG:00190), mitochondrial complex I assembly (WP:4324), electron transport chain (WP:111), and mitochondrial complex IV assembly (WP:4922), together with a parallel upregulation of the calcium signaling pathway (KEGG:04020). To determine whether the mitochondrial transcriptional signature identified at the aggregate DEG level (Figure 1B) is driven by cell-type-specific changes in inhibitory neurons versus excitatory neurons, we performed cell-type-resolved reanalysis of the same scRNA-seq dataset with cells annotated by lineage (Figures S1A). We then examined expression of five canonical regulators of mitochondrial biogenesis and dynamics, *PPARGC1A* (PGC1α) and *TFAM* (biogenesis), *DNM1L* (DRP1) and *FIS1* (fission), and *OPA1* (fusion), in excitatory versus inhibitory neurons (Figure S1B and S1C). Again, the inhibitory lineage showed the most consistent and pronounced dysregulation. 22q11.2DS inhibitory neurons exhibited significant downregulation of all five mitochondrial regulators (Wilcoxon rank-sum test; PPARGC1A p = 5.6 × 10⁻[; TFAM p = 3.2 × 10⁻¹□; DNM1L p = 1.1 × 10⁻¹□; OPA1 p = 1.5 × 10⁻□), with FIS1 uniquely unchanged (p = 0.907). Statistical significance was determined by the Wilcoxon rank-sum test for all comparisons.

Together, our new analyses of the published dataset suggest that (i) genes associated with neuropsychiatric diseases are preferentially differentially expressed in inhibitory neurons in 22q11.2DS, and that (ii) mitochondrial dysfunction is enriched in inhibitory neuronal lineages in 22q11.2DS, thus supporting our scientific endeavor in this manuscript.

### 22q11.2DS Patient-Derived Subpallial Organoids (hSOs) Exhibit Transcriptomic Alterations

To investigate the cellular consequences of 22q11.2 deletion in inhibitory interneurons, we generated human subpallial organoids (hSOs) from four control (2242-1, 1205-4, 1208-2, 8119-1) and four 22q11.2DS patient-derived induced pluripotent stem cell (iPSC) lines (6303-5, 7958-3, 1804-5, 9050-3) (Figure 1C). Using bulk RNA-seq in control and 22q11.2DS hSOs from 4 lines each at day 100 in culture, we confirmed hemizygosity of the 22q11.2 locus for gene expression (Figure 1D).

While bulk RNAseq did not suggest any transcriptional differences in the progression of maturation from progenitors to neurons or in the subtypes of generated interneurons at day 100 (Figure S2), we independently quantified the proportions of SOX2+ neural progenitors and TUJ1+ neurons at day 100 using flow cytometry, as well as the transcriptional expression of FOXG1 as a dorsal forebrain marker, and that of markers associated with various subtypes of interneurons (somatostatin (*SST*), calbindin (*CALB*), parvalbumin (*PVALB*)). Quantification of these markers revealed comparable proportions of TUJ1+ neurons (p = 0.2141; Welch’s t-test, Figure S2A) and SOX2+ progenitors (p = 0.2044; Welch’s t-test, Figure S2A), no transcriptional differences of other markers (Figure S2B), and high sample correlation (Figure S2C), suggesting no major differentiation defects.

For analyses, we first repeated the overlap analysis of 22q11.2DS differentially expressed genes (DEGs) in our dataset from hSOs, with published neuropsychiatric disorder risk gene sets; we demonstrated similar convergence with risk genes for schizophrenia (SCZ), autism spectrum disorder (ASD), major depressive disorder (MDD), bipolar disorder (BD), and ADHD, with both upregulated and downregulated DEGs contributing to these overlaps (Figure 1E).

Among the upregulated pathways, we found the glycolytic process (NES = +1.32) and cellular response to hypoxia (NES = +1.53) to be prominently enriched, indicating a metabolic shift toward glycolysis and decreased mitochondrial respiration in 22q11.2DS interneurons; interestingly, these processes are seen in hypoxia-exposed control organoids^20,25,26^. Pathways related to mitochondrial fragmentation (NES = +0.60), regulation of mitochondrial fission (NES = +0.45), and negative regulation of mitochondrial fusion (NES = +0.44) were also consistently upregulated, suggesting a coordinated shift toward mitochondrial fragmentation, a morphological marker of stress. Actin crosslink formation (NES = +0.86) and inflammatory response pathways (NES = +0.97) were also enriched among upregulated genes (Figure 1F). Conversely, pathways essential for neuronal function and mitochondrial energy production were significantly downregulated. Neurotransmitter secretion (NES =-2.08) and regulation of synapse assembly (NES =-1.52) were among the most depleted, consistent with impaired synaptic maturation. Mitochondrial ATP synthesis coupled to electron transport (NES =-1.56) and oxidative phosphorylation (NES =-1.08) were both downregulated, reinforcing the interpretation of a glycolysis/OXPHOS metabolic imbalance. Additional downregulated pathways included mitochondrion distribution (NES =-1.46, axon development (NES =-1.20), voltage-gated calcium channel activity (NES =-0.93), inner mitochondrial membrane organization (NES =-0.80), regulation of kinase activity (NES =-0.62), and store-operated calcium entry (NES =-0.48) (Figure 1F). Statistical significance was determined by Fisher’s exact test and GSEA pre-ranked analysis.

A separate categorization of DEGs by mitochondrial functional class - done using mitoXplorer3.0^25^- further corroborated these findings (Figure 1G). Glycolysis exhibited the greatest imbalance, with 13 upregulated and only 2 downregulated genes, while oxidative phosphorylation showed the inverse pattern (14 downregulated, 2 upregulated). Additional categories with notable dysregulation included mitochondrial dynamics, calcium signaling and transport, and reactive oxygen species defense. Specifically, glycolytic genes including *LDHA*, *HK2*, *PKM*, *ENO1*, *SLC2A1*, *GAPDH*, *PFKP*, *TPI1*, *PGAM1*, *GPI*, *PC*, and *PGK1* were upregulated, while OXPHOS genes such as *NDUFA8*, *NDUFA6*, *COX20*, *ATP5PB*, *COX5A*, *ATP5MC1*, *UQCRC2*, *NDUFA1*, *ETFDH*, and *HIGD1A* were downregulated (Figure S2D).

These differences clearly show a global and significant effect of the deletion on overall mitochondrial function and, interestingly, suggest potential phenotypic overlap between 22q11.2DS and hypoxic injuries.

Given the strong signal in mitochondrial function-related genes, we separately examined changes in their transcriptional expression within the 22q11.2 hemizygously deleted region. The 22q11.2 region-specific genes, including *PRODH*, *TANGO2*, *MRPL40*, *ZDHHC8*, *TXNRD2*, *SLC25A1*, *COMT*, and *DGCR8*, showed expected reductions in expression consistent with the expected effects of hemizygosity (Figure S2E).

Overall, these data reveal a convergent transcriptomic signature in 22q11.2DS hSOs, characterized by a shift from oxidative phosphorylation toward glycolysis, increased mitochondrial fragmentation, and impaired synaptic and calcium signaling pathways.

### 22q11.2DS Patient-Derived Subpallial Organoids Display Mitochondrial Respiration and Morphology Defects

To determine whether the transcriptomic metabolic shift in 22q11.2DS hSOs corresponds to functional mitochondrial impairments, we performed Seahorse XF Mito Stress test on dissociated hSOs at day 100, from 4 control and 4 22q11.2DS lines. hSOs showed significant reductions in all measured respiration parameters compared to controls: basal respiration (p = 0.0002; Welch’s t-test), maximal respiration (p = 0.0002; Welch’s t-test), spare respiratory capacity (p = 0.0006; Welch’s t-test), non-mitochondrial respiration (p = 0.0047; Welch’s t-test), ATP production (p = 0.0002; Welch’s t-test), and proton leak (p = 0.0104; Welch’s t-test) (Figures 2A and 2B). These results confirm findings from transcriptomics, suggesting a genuine deficit in mitochondrial oxidative capacity in hSOs.

**Figure 2.**
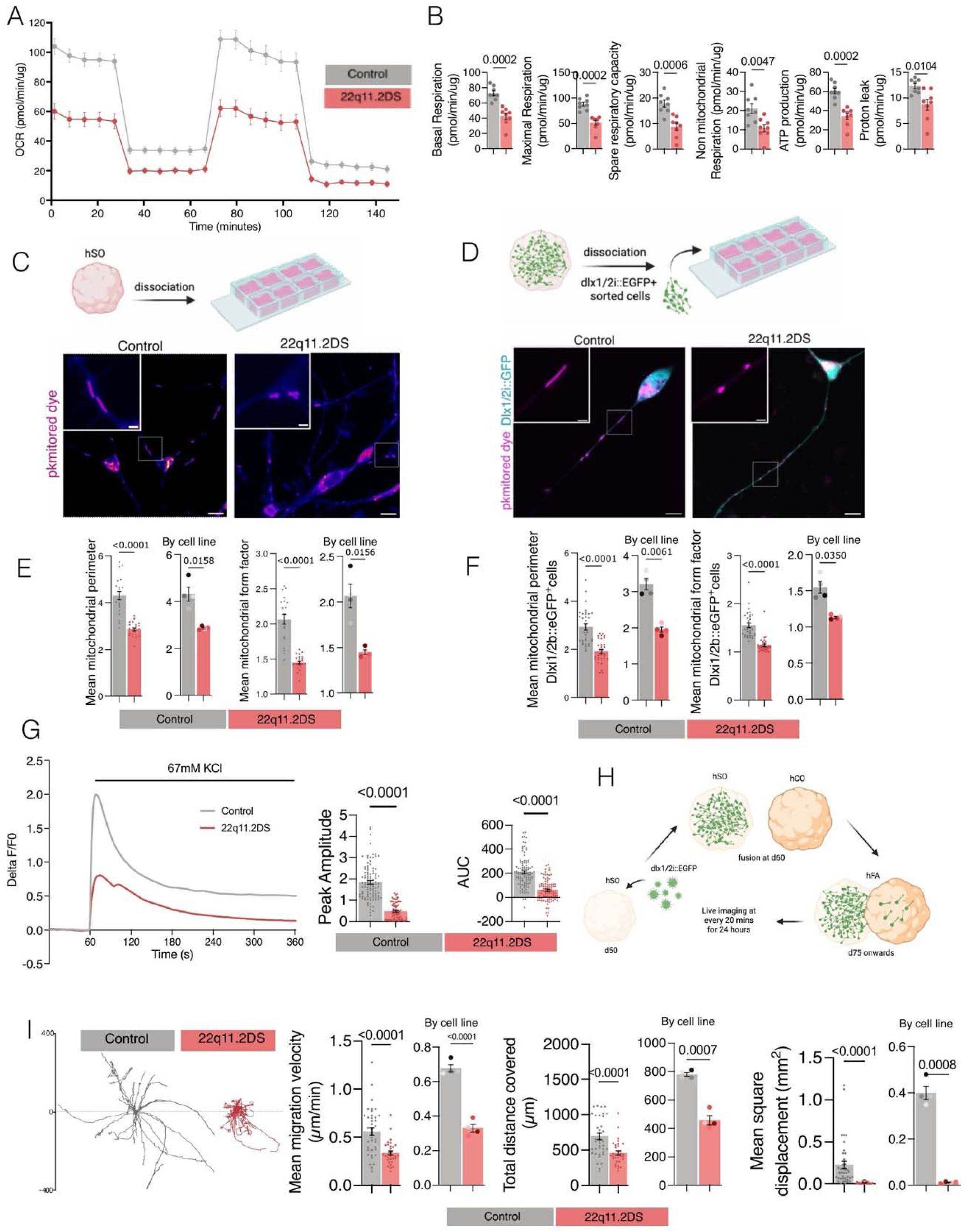
Mitochondrial and interneuron migration defects in 22q11.2DS models 22q11.2DS hSOs show major deficits in mitochondrial respiration and structure, along with reduced interneuron migration speed, distance, and displacement in human forebrain assembloids (hFAs). **A)** Seahorse XF Mito Stress Test. Oxygen consumption rate (OCR; pmol/min/µg) over time in control (gray) and 22q11.2DS (red) hSOs. Sequential injections of oligomycin, FCCP, and rotenone/antimycin A are marked. **(B)** Respiration parameter measurements: basal respiration (p = 0.0002, Welch’s t-test), maximal respiration (p = 0.0002, Welch’s t-test), spare respiratory capacity (p = 0.0006, Welch’s t-test), non-mitochondrial respiration (p = 0.0047, Welch’s t-test), ATP production (p = 0.0002, Welch’s t-test), and proton leak (p = 0.0104, Welch’s t-test). Each dot shows one organoid; n = 8 (control) and 8 (22q11.2DS) organoids from 4 iPSC lines per genotype. **(C)** Top: diagram of hSO dissociation for mitochondrial shape analysis. Bottom: representative confocal images of mitochondria stained with pkMitoDeepRed in dissociated bulk hSO neurons. Control vs. 22q11.2DS. Insets show enlarged mitochondria. Scale bar, 20 µm. **(D)** Top: diagram of Dlx1/2i::EGFP+ cell sorting. Bottom: representative confocal images of mitochondrial shape (pkMitoDeepRed, magenta) in sorted Dlx1/2i::eGFP+ interneurons (cyan). Control vs. 22q11.2DS. Insets show magnified mitochondria. Scale bar, 20 µm. **(E)** Mitochondrial shape analysis in bulk hSO neurons: average mitochondrial perimeter (p < 0.0001, p = 0.0158, Welch’s t-test) and average mitochondrial form factor (p < 0.0001, p = 0.0156, Welch’s t-test). Each dot is a cell; n = 21 (control) and 22 (22q11.2DS) cells from 4 iPSC lines per genotype. **(F)** Mitochondrial shape analysis in Dlx1/2i::eGFP+ interneurons: average mitochondrial perimeter (p < 0.0001, p = 0.0061, Welch’s t-test) and average mitochondrial form factor (p < 0.0001, p = 0.0350, Welch’s t-test). Each dot is a cell; n = 33 (control) and 32 (22q11.2DS) cells from 4 iPSC lines per genotype. **(G)** Left: Calcium transient traces (ΔF/F₀) upon 67 mM KCl stimulation in control (gray) and 22q11.2DS (red) hSOs over 360 s. Right: quantification of peak amplitude (p < 0.0001, Welch’s t-test) and area under the curve (AUC; p < 0.0001, Welch’s t-test). Each dot represents one neuron; n = 94 (control) and 75 (22q11.2DS) neurons from 4 iPSC lines per genotype. **(H)** Diagram of the human forebrain assembloid (hFA) migration test. hSOs expressing Dlx1/2i::EGFP were combined with human cortical organoids (hCOs) at day 60 (d60). Live imaging was performed every 20 min over 24 h, starting from d75. **(I)** Left: migration paths of Dlx1/2i::EGFP+ interneurons plotted from a common origin over ±400 µm. Control (gray) vs. 22q11.2DS (red). Right: Migration analysis. Average migration speed (µm/min; p < 0.0001, Welch’s t-test), total distance traveled (µm; p < 0.0001, p = 0.0007, Welch’s t-test) and mean square displacement (mm²; p < 0.0001, p = 0.0008, Welch’s t-test). Each dot is a cell; n = 39 (control) and 33 (22q11.2DS) cells from 4 iPSC lines per genotype. Data are shown as mean ± SEM. See also Figures S2 and S3.

Given that mitochondrial stress is often associated with morphologic changes, we next assessed mitochondrial morphology in control and 22q11.2DS hSOs using pkMitoDeepRed staining. Confocal imaging demonstrated significant mitochondrial fragmentation in 22q11.2DS. In dissociated bulk hSO neurons, 22q11.2DS cells had markedly reduced mean mitochondrial perimeter (p < 0.0001; Welch’s t-test) and form factor (p < 0.0001; Welch’s t-test), indicating smaller, more circular (fragmented) mitochondria (Figures 2C-F). Although hSOs contain almost exclusively cortical interneurons and ventral progenitors lineage, we further increased the robustness and specificity of the observed phenotype by analyzing FACS-sorted Dlx1/2i::EGFP+ interneurons, which similarly exhibited lower perimeter (p < 0.0001) and form factor (p < 0.0001) (Figures 2D and 2F). Importantly, qRT-PCR analysis of *DLX1* (p = 0.7623; Welch’s t-test) and *GAD2* (p = 0.5254; Welch’s t-test) in sorted Dlx1/2i::GFP+ cells showed comparable expression of interneuron markers between genotypes, ruling out maturation stage and cell fate of the Dlx1/2i::EGFP+ interneuron as a confounding factor for the observed phenotype (Figure S3A). Details on number of samples are presented in Table S1.

### 22q11.2DS Patient-Derived Subpallial Organoids Exhibit Calcium Signaling Defects

Based on results from bulk RNA-seq in hSOs, which suggested abnormal calcium biology, we functionally probed calcium signaling in control versus 22q11.2DS hSOs derived from 4 lines per phenotype at 100 days. We observed significant calcium-signaling deficits in monolayer cultures derived from dissociated 22q11.2DS hSOs. Upon 67 mM KCl stimulation, Calbryte-520-AM-loaded neurons exhibited reduced peak amplitude (p < 0.0001; Welch’s t-test) and area under the curve (AUC; p < 0.0001; Welch’s t-test) compared to controls (Figure 2G). These newly identified defects in inhibitory interneurons align with previous reports from excitatory neurons carrying the 22q11.2 deletion, suggesting this phenotype is shared among multiple neuronal subtypes.

### Impaired Cortical Interneuron Migration in 22q11.2DS

To assess interneuron migration in 22q11.2DS, we generated human forebrain assembloids (hFAs) by fusing Dlx1/2i::EGFP-labeled hSOs with human cortical organoids (hCOs) at ∼day 60 of differentiation (d60), using previously published protocols^20, 21^. Two weeks later, we performed live imaging on interneurons migrated to the hCO side; images were obtained every 20 minutes for 24 (Figure 2H). Migration tracking of Dlx1/2i::EGFP+ interneurons on the hCOs side showed that 22q11.2DS cells traveled shorter distances with less displacement from their origin compared to controls. Quantification confirmed significant decreases in mean migration velocity (µm/min; p < 0.0001; Welch’s t-test), total distance (µm; p < 0.0001; Welch’s t-test), and mean square displacement (mm²; p < 0.0001; Welch’s t-test), a measure of directed motility, in 22q11.2DS hFA (Figure 2I). Together, these findings demonstrate a combined mitochondrial and migration deficits in 22q11.2DS interneurons. Statistical analysis used Welch’s t-test (B, E, G, and I). Details on number of samples are presented in Table S1.

### Adrenomedullin (ADM) Rescues Mitochondrial and Migration Deficits in 22q11.2DS Interneurons

ADM^27^ is a 52-amino-acid peptide with an intramolecular disulfide bridge and an amidated C-terminus that signals through the calcitonin receptor-like receptor (CLR)/receptor activity-modifying protein 2 (RAMP2) complex, engaging protein kinase A^28^ (PKA; via Gα → adenylyl cyclase [AC] → cyclic AMP [cAMP] → PKA) and phospholipase C^29^ (PLC; via Gα → PLC → inositol trisphosphate [IP₃] → Ca²⁺) signaling cascades (Figure 3A). A previous report from our laboratory showed that exogenous ADM supplementation rescued hypoxic interneuron migration deficits^20^. Since both hypoxia and 22q11.2 display mitochondrial dysfunction, we tested whether ADM could rescue the migration of 22q11.2DS interneurons and their associated mitochondrial phenotypes.

**Figure 3.**
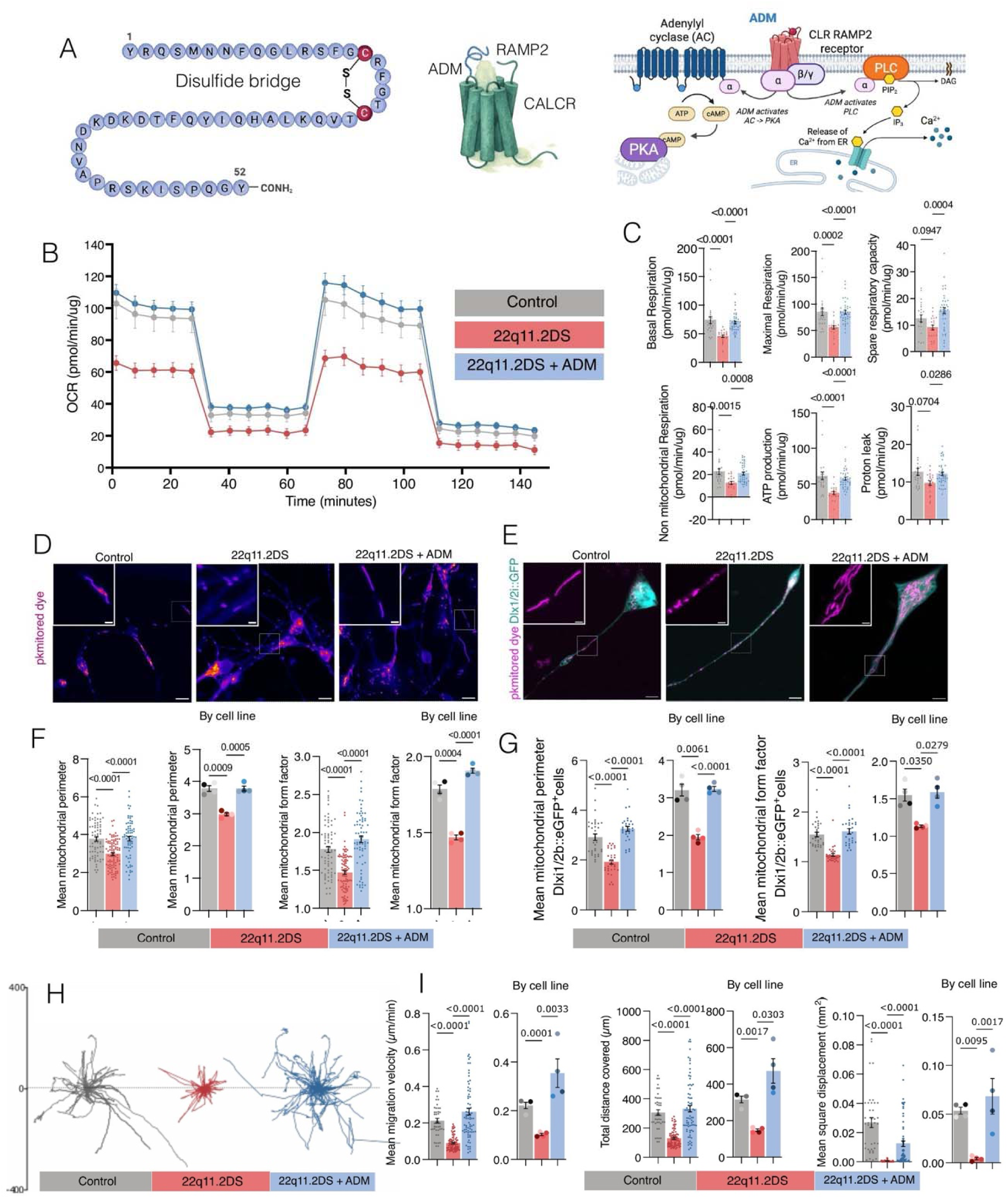
Adrenomedullin Rescues Interneuron Migration and Mitochondrial Deficits in 22q11.2DS Cellular Models. Adrenomedullin (ADM) treatment rescues mitochondrial respiration, morphology, and interneuron migration in 22q11.2DS hSOs and forebrain assembloids. **(A)** Left: amino acid sequence and structure of adrenomedullin (ADM; 52 amino acids) with an intramolecular disulfide bridge and amidated C-terminus (CONH₂). Right: schematic of ADM signaling through the calcitonin receptor-like receptor (CLR)/receptor activity-modifying protein 2 (RAMP2) complex. ADM binding activates two downstream pathways: protein kinase A (PKA; Gα subunit of ADM-bound receptor → adenylyl cyclase [AC] → ATP → cyclic AMP [cAMP] → PKA) and phospholipase C (PLC; Gα subunit of ADM-bound receptor → PLC → phosphatidylinositol 4,5-bisphosphate [PIP₂] → diacylglycerol [DAG] + inositol trisphosphate [IP₃] → release of Ca²⁺ from endoplasmic reticulum [ER]). **(B)** Seahorse OCR traces (pmol/min/µg) over time in control (gray), 22q11.2DS (red), and 22q11.2DS + ADM (blue) hSOs. Sequential injections of oligomycin, FCCP, and rotenone are indicated. **(C)** Quantification of respiration parameters across three conditions: basal respiration (p < 0.0001, p < 0.0001, Kruskal-Wallis test with Dunn’s post hoc multiple comparisons), maximal respiration (p = 0.0001, p = 0.0004, Kruskal-Wallis test with Dunn’s post hoc multiple comparisons), spare respiratory capacity (p < 0.0001, Kruskal-Wallis test with Dunn’s post hoc multiple comparisons), non-mitochondrial respiration (p = 0.0008, p = 0.0015, Kruskal-Wallis test with Dunn’s post hoc multiple comparisons), ATP production (p < 0.0001, Kruskal-Wallis test with Dunn’s post hoc multiple comparisons), and proton leak (p = 0.0704, p = 0.0286, Kruskal-Wallis test with Dunn’s post hoc multiple comparisons). Each dot represents one organoid; n = 22 (control), 20 (22q11.2DS), and 40 (22q11.2DS + ADM) organoids from 4 iPSC lines per condition. **(D)** Representative confocal images of mitochondrial morphology (pkMitoDeepRed) in bulk hSO neurons across three conditions: control, 22q11.2DS, and 22q11.2DS + ADM. Insets show magnified mitochondria. Scale bar, 20 µm. **(E)** Representative confocal images of mitochondrial morphology (pkMitoDeepRed, magenta) in Dlx1/2i::eGFP+ sorted interneurons (cyan) across three conditions. Scale bar, 20 µm. **(F)** Quantification of mitochondrial morphology in bulk neurons: mean mitochondrial perimeter (p < 0.0001, p = 0.0005, p < 0.0001, Kruskal-Wallis test with Dunn’s post hoc multiple comparisons) and mean form factor (p < 0.0001, p = 0.0098, Kruskal-Wallis test with Dunn’s post hoc multiple comparisons). Each dot represents one cell; n = 74 (control), 82 (22q11.2DS), and 70 (22q11.2DS + ADM) cells from 4 iPSC lines per condition. **(G)** Quantification of mitochondrial morphology in Dlx1/2i::eGFP+ interneurons: mean mitochondrial perimeter (p < 0.0001, p = 0.0061, p < 0.0001, Kruskal-Wallis test with Dunn’s post hoc multiple comparisons) and mean form factor (p < 0.0001, p = 0.0350, p = 0.0275, Kruskal-Wallis test with Dunn’s post hoc multiple comparisons). Each dot represents one cell; n = 33 (control), 32 (22q11.2DS), and 30 (22q11.2DS + ADM) cells from 4 iPSC lines per condition. **(H)** Representative migration tracks of Dlx1/2i::EGFP+ interneurons in hFAs plotted from a common origin over ±400 µm: control (gray), 22q11.2DS (red), and 22q11.2DS + ADM (blue). **(I)** Quantification of migration: mean migration velocity (µm/min; p < 0.0001, p = 0.0023, Kruskal-Wallis test with Dunn’s post hoc multiple comparisons), total distance covered (µm; p < 0.0001, p = 0.0017, Kruskal-Wallis test with Dunn’s post hoc multiple comparisons) and mean square displacement (mm²; p < 0.0001, p = 0.0095, p = 0.0017, Kruskal-Wallis test with Dunn’s post hoc multiple comparisons). Each dot represents one tracked cell; n = 44 (control), 77 (22q11.2DS), and 75 (22q11.2DS + ADM) cells from 4 iPSC lines per condition. Data are represented as mean ± SEM. See also Figure S4.

First, we checked for differences in baseline ADM expression levels between 22q11.2DS and control hSOs. qRT-PCR analysis confirmed that *ADM* (p = 0.4428) and its receptor components *CRLR* (p = 0.9893), *RAMP1* (p = 0.9323), *RAMP2* (p = 0.7987), and *RAMP3* (p = 0.3697) are comparably expressed between genotypes, indicating that the ADM signaling machinery is intact in 22q11.2DS hSOs (Figure S4A).

Upon ADM supplementation (0.5 µM) of 22q11.2DS hSOs for 24 hours, we observed restoration of all major phenotypes, as described below.

Seahorse analysis of three conditions (control, 22q11.2DS, and 22q11.2DS + ADM) demonstrated restoration of basal respiration (p < 0.0001), maximal respiration (p < 0.0001), spare respiratory capacity (p = 0.0004), non-mitochondrial respiration (p = 0.0015), ATP production (p < 0.0001), and proton leak (p = 0.0286) (Figures 3B and 3C).

ADM supplementation also reversed the mitochondrial fragmentation phenotype. Confocal imaging of pkMitoDeepRed-stained mitochondria demonstrated restored elongated morphology in 22q11.2DS + ADM bulk neurons and interneurons compared to untreated 22q11.2DS cells (Figures 3D and 3E). Quantification in bulk neurons showed significant rescue of mean mitochondrial perimeter (p < 0.0001 for control vs. 22q11.2DS; p = 0.0005 for 22q11.2DS vs. 22q11.2DS + ADM; p < 0.0001 for control vs. 22q11.2DS + ADM) and form factor (p < 0.0001; p = 0.8004) (Figure 3F). This rescue was further confirmed in sorted Dlx1/2i::eGFP+ interneurons, where ADM rescued mitochondrial perimeter (p < 0.0001, p = 0.0061, p < 0.0001) and form factor (p < 0.0001, p = 0.0350, p = 0.0279) (Figure 3G).

Critically, ADM supplementation also restored interneuron migration in hFAs. Migration tracks of 22q11.2DS + ADM interneurons (blue) showed recovery toward control (gray) patterns, unlike untreated 22q11.2DS cells (red) (Figure 3H). Quantification revealed significant rescue of mean velocity (µm/min; p < 0.0001, p = 0.0033), saltation (p < 0.0001, p = 0.0001), total distance (µm; p < 0.0001, p = 0.0303), and mean square displacement (mm²; p < 0.0001, p = 0.0017) (Figure 3I). These results identify ADM as a powerful rescue agent capable of restoring both mitochondrial function and interneuron migration in 22q11.2DS cellular models. Statistical significance was determined by the Kruskal-Wallis test with Dunn’s post hoc multiple comparisons (C, F, G, and I). Details on the number of samples are presented in Table S1.

### Mitochondrial and Migration Deficits Are Recapitulated in Primary *ex vivo* Human Cortical Tissue and Rescued by ADM

To determine whether the phenotypes seen in organoid models reflect bona fide features of 22q11.2DS in developing human brain tissue, we obtained primary ex vivo human cortical tissue (hCT) at gestational ages of 19 weeks 2 days (19w2d; control) and 19 weeks 4 days (19w4d; 22q11.2DS). The brain tissue was sectioned into 2 mm slices, infected with Dlx1/2i::EGFP lentivirus, and subjected to live imaging every 20 minutes for 24 hours (Figure 4A).

**Figure 4.**
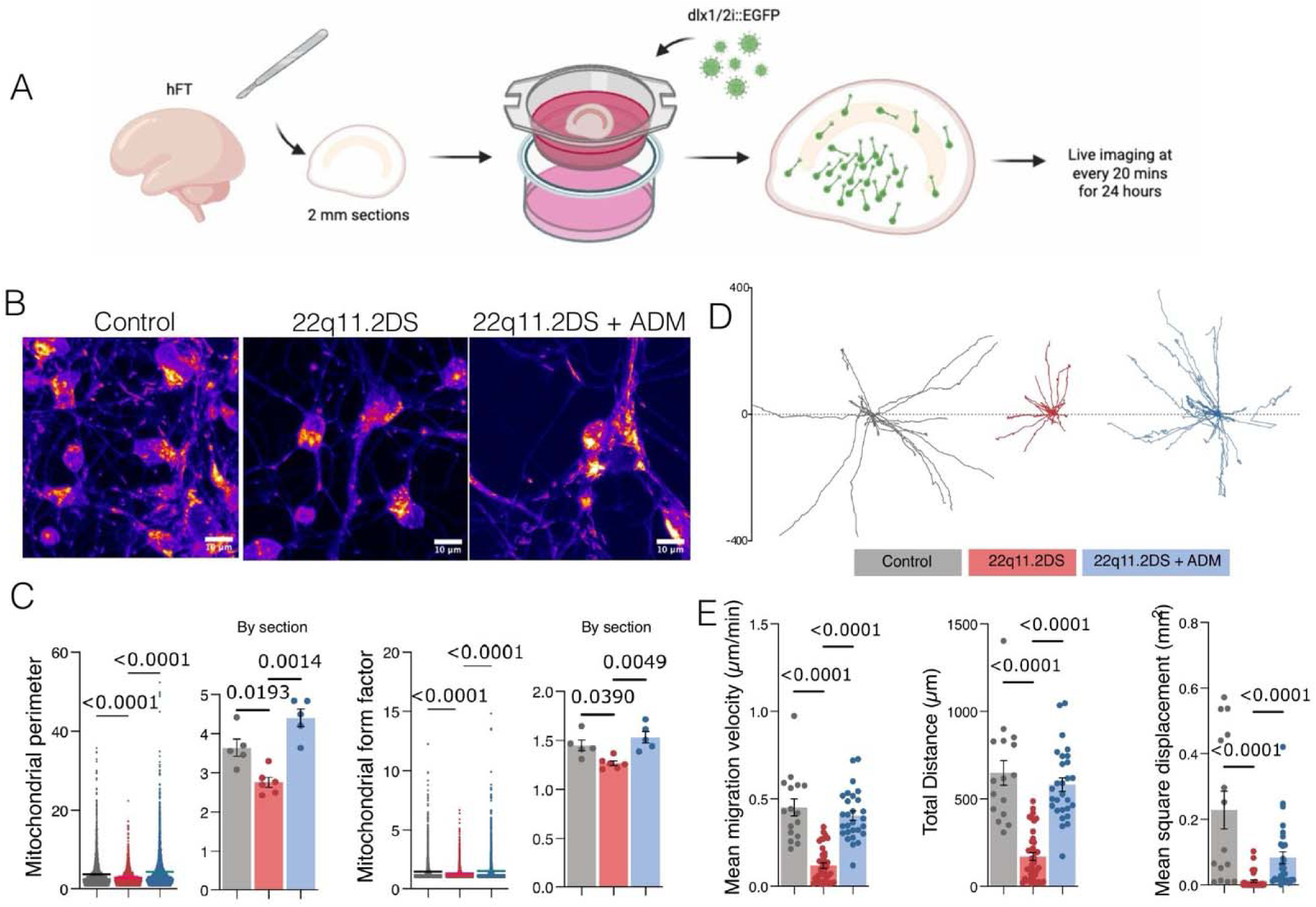
Validation of Mitochondrial and Migration Deficits in Human Cortical Tissue with 22q11.2 Deletion and Rescue with Adrenomedullin. Mitochondrial morphology defects and impairments in interneuron migration observed in hSOs are recapitulated in human cortical tissue (hCT) carrying the 22q11.2 deletion and are rescued by ADM treatment. **(A)** Schematic of experimental design. Human cortical tissue (hCT) was sectioned into 2 mm slices, infected with Dlx1/2i::EGFP lentivirus, and live-imaged every 20 min for 24 h. **(B)** Representative confocal images of mitochondria in hCT: control 19w2d, 22q11.2DS 19w4d, and 22q11.2DS + ADM. A hot look-up table (LUT; yellow magenta) is shown; structural staining is in blue. Scale bar, 20 µm. **(C)** Quantification of mitochondrial perimeter (p < 0.0001, p = 0.0193, p = 0.0014, Kruskal-Wallis test with Dunn’s post hoc multiple comparisons) and form factor (p < 0.0001, p = 0.0049, p = 0.0390, Kruskal-Wallis test with Dunn’s post hoc multiple comparisons). Each dot represents one cell; n = 3,162 (control), 2,341 (22q11.2DS), and 3,085 (22q11.2DS + ADM) cells from 5, 6, and 5 technical replicates per condition, respectively. **(D)** Representative migration tracks: control (gray), 22q11.2DS (red), and 22q11.2DS + ADM (blue). **(E)** Quantification of migration: mean velocity (µm/min; p < 0.0001, Kruskal-Wallis test with Dunn’s post hoc multiple comparisons), total distance (µm; p < 0.0001, Kruskal-Wallis test with Dunn’s post hoc multiple comparisons), and mean square displacement (mm²; p < 0.0001, Kruskal-Wallis test with Dunn’s post hoc multiple comparisons). Each dot represents one tracked cell; n = 16 (control), 39 (22q11.2DS), and 28 (22q11.2DS + ADM) cells from 3 biological replicates per condition. Data are represented as mean ± SEM. The n values are as indicated per panel from 3 biological replicates.

Confocal imaging of mitochondria in hCT revealed the same fragmentation phenotype observed in organoids: 22q11.2DS hCT had shorter, more circular mitochondria compared to control tissue (Figure 4B); ADM supplementation restored mitochondrial morphology in 22q11.2DS hCT. Quantification confirmed significant reductions in mitochondrial perimeter (p < 0.0001 for control vs. 22q11.2DS) and form factor (p < 0.0001) in 22q11.2DS tissue, with ADM rescuing both perimeter (p = 0.0193 for 22q11.2DS vs. 22q11.2DS + ADM; p = 0.0014 for control vs. 22q11.2DS + ADM) and form factor (p = 0.0049; p = 0.0390) (Figure 4C).

Interneuron migration was similarly impaired in 22q11.2DS hCT and rescued by ADM. Specifically, migration tracks showed reduced displacement in 22q11.2DS interneurons compared with controls, with recovery upon ADM supplementation at 0.5 µM for 24 hours (Figure 4D). Quantification of mean velocity (µm/min; p < 0.0001), total distance (µm; p < 0.0001), and mean square displacement (mm²; p < 0.0001) confirmed significant impairment across all metrics in 22q11.2DS hCT, with ADM restoring all parameters toward control levels (Figure 4E).

This represents the first demonstration that 22q11.2DS mitochondrial and migration phenotypes exist in primary human cortical tissue, and that ADM can rescue them ex vivo, thus establishing the translational relevance of our organoid findings. Statistical significance was determined by the Kruskal-Wallis test with Dunn’s post hoc multiple comparisons (C and E). Details on number of samples are presented in Table S1.

### ADM Restores Mitochondrial Membrane Potential, DRP1 Phosphorylation, Calcium Homeostasis, and Actin Dynamics

To elucidate the mechanism by which ADM rescues mitochondrial and interneuron migration phenotypes, we first examined DRP1 phosphorylation, a key regulatory node in mitochondrial fission-fusion dynamics (Figure 5A). Phosphorylation at Ser616 promotes mitochondrial fission, while phosphorylation at Ser637 inhibits DRP1 and promotes fusion. Western blot analysis of control, 22q11.2DS, and 22q11.2DS + ADM hSOs showed that 22q11.2DS organoids have higher pSer616/DRP1 ratios and lower pSer637/DRP1 ratios than controls, consistent with a pro-fission state (Figures 5B and 5C). ADM treatment significantly lowered the pSer616/DRP1 ratio (p < 0.0001, p = 0.0047) and increased the pSer637/DRP1 ratio (p = 0.0034, p = 0.0470), shifting the balance toward mitochondrial fusion (Figure 5C).

**Figure 5.**
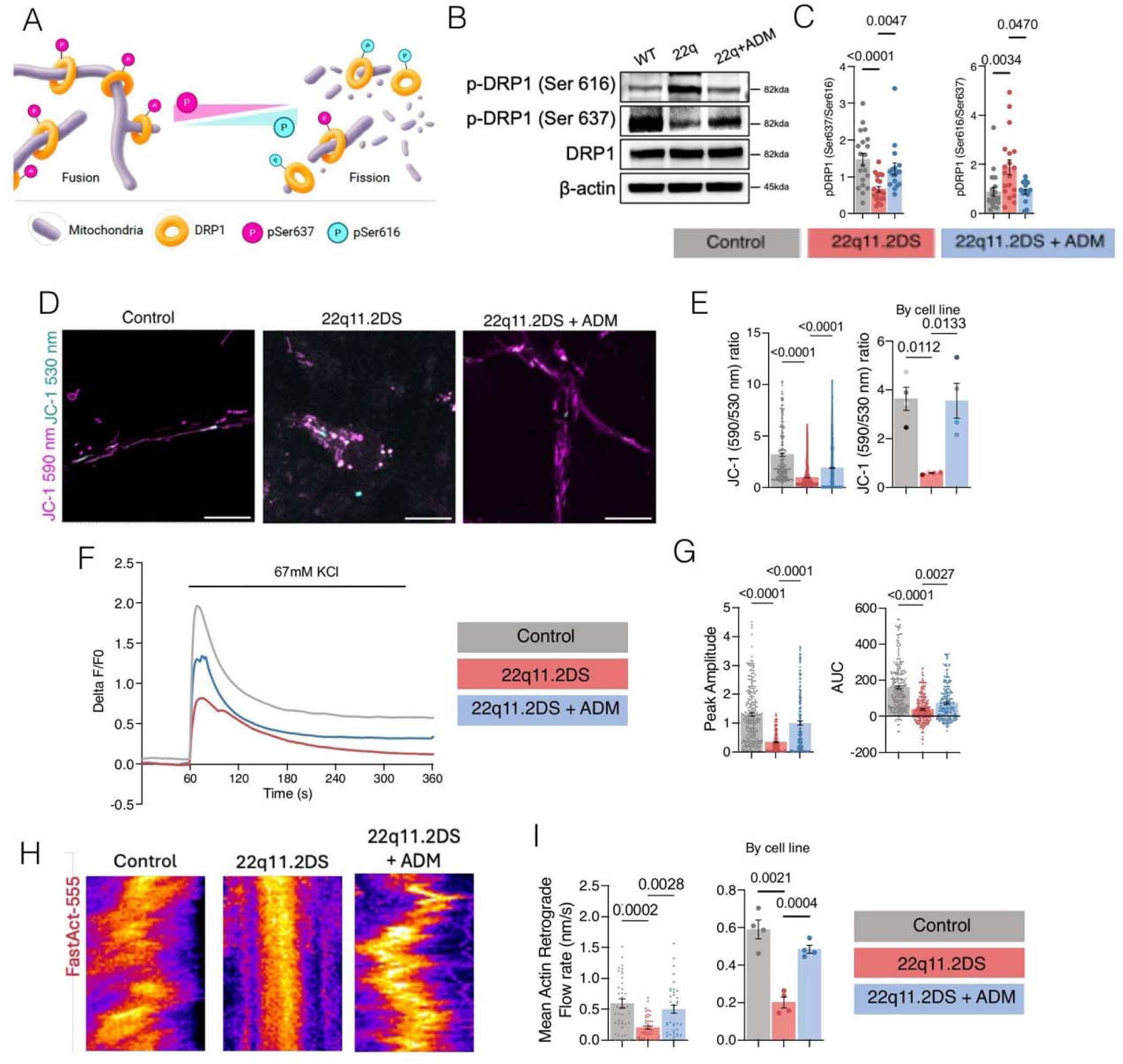
Mechanistic Insights: ADM Restores Mitochondrial Function, Calcium Homeostasis, and Cytoskeletal Dynamics. ADM shifts DRP1 phosphorylation toward the pro-fusion site Ser637, restores mitochondrial membrane potential (MMP), rescues calcium transients, and normalizes actin retrograde flow in 22q11.2DS interneurons. **(A)** Schematic of dynamin-related protein 1 (DRP1)-mediated mitochondrial fission and fusion balance. Phosphorylation at Ser637 (yellow) promotes fusion; phosphorylation at Ser616 (red) promotes fission. **(B)** Western blot analysis of phospho-DRP1 (Ser616), phospho-DRP1 (Ser637), total DRP1, and β-actin in control, 22q11.2DS, and 22q11.2DS + ADM hSOs. Molecular weights: 82 kDa (DRP1), 45 kDa (β-actin). **(C)** Quantification of pSer616/DRP1 (p < 0.0001, p = 0.0047, Kruskal-Wallis test with Dunn’s post hoc multiple comparisons) and pSer637/DRP1 (p = 0.0034, p = 0.0470, Kruskal-Wallis test with Dunn’s post hoc multiple comparisons) ratios. Each dot represents one biological replicate: n = 19 (control), 19 (22q11.2DS), and 16 (22q11.2DS + ADM). **(D)** Representative JC-1 mitochondrial membrane potential (MMP) images: JC-1 590 nm (magenta, J-aggregates indicating high MMP) and JC-1 530 nm (cyan, monomers indicating low MMP) in control, 22q11.2DS, and 22q11.2DS + ADM. Scale bar, 20 µm. **(E)** JC-1 590/530 nm ratio quantified by cell (p < 0.0001, Kruskal-Wallis test with Dunn’s post hoc multiple comparisons) and by replicate (p = 0.0133, p = 0.0112, Kruskal-Wallis test with Dunn’s post hoc multiple comparisons). Each dot represents one cell or one replicate as indicated; n = 250 (control), 1,362 (22q11.2DS), and 1,874 (22q11.2DS + ADM) cells from 4 iPSC lines per condition. **(F)** Calcium transient traces (ΔF/F₀, 67 mM KCl stimulation) in control (gray), 22q11.2DS (red), and 22q11.2DS + ADM (blue) hSOs over 360 s. **(G)** Quantification of peak amplitude (p < 0.0001, Kruskal-Wallis test with Dunn’s post hoc multiple comparisons) and AUC (p = 0.0027, p < 0.0001, Kruskal-Wallis test with Dunn’s post hoc multiple comparisons). Each dot represents one neuron; n = 241 (control), 209 (22q11.2DS), and 212 (22q11.2DS + ADM) neurons from 4 iPSC lines per condition. **(H)** Representative kymographs of actin retrograde flow (FastAct-555) in growth cones: control, 22q11.2DS, and 22q11.2DS + ADM. Scale bar, 20 µm. **(I)** Mean actin retrograde flow rate (p = 0.0028, p = 0.0004, Kruskal-Wallis test with Dunn’s post hoc multiple comparisons) and additional flow parameters as shown. Each dot represents a unique kymograph per growth cone; n = 43 (control), 47 (22q11.2DS), and 37 (22q11.2DS + ADM) growth cones from 4 iPSC lines per condition. Data are shown as mean ± SEM. n values are as indicated per panel from 4 iPSC lines per condition.

Since DRP1 phosphorylation balance is directly related to the mitochondrial membrane potential (MMP), we next assessed this in hSOs using the ratiometric dye JC-1. This analysis revealed that 22q11.2DS hSOs exhibit decreased MMP, as evidenced by a lower 590/530 nm ratio, reflecting a shift from J-aggregates (high MMP, 590 nm) toward monomers (low MMP, 530 nm). This decrease in MMP is associated with fragmented mitochondria. However, this was restored by ADM supplementation (Figures 5D and 5E), both at the single-cell level (p < 0.0001) and across biological replicates (p = 0.0133, p = 0.0112) (Figure 5E).

Next, we checked whether ADM also rescues the observed calcium signaling deficits. Calcium imaging upon 67 mM KCl stimulation showed that 22q11.2DS + ADM hSOs exhibited restored calcium transients (blue) approaching control levels (gray), in contrast to the diminished responses in untreated 22q11.2DS cells (red) (Figure 5F). Quantification demonstrated significant rescue of peak amplitude (p < 0.0001) and AUC (p = 0.0027, p < 0.0001) (Figure 5G).

Given the critical roles of mitochondrial energy production and calcium signaling in actomyosin contractility - a key feature of neuronal migration, we examined the F-actin retrograde flow in interneuron growth cones. Analysis of actin retrograde flow in neuronal growth cones using FastAct-555 labeling showed that 22q11.2DS interneurons have reduced flow rates compared to controls, but this was restored with ADM supplementation (Figures 5H and 5I). Quantification confirmed a significant rescue of the mean flow rate (p = 0.0028, p = 0.0004) (Figure 5I). Statistical significance was assessed using Kruskal-Wallis test with Dunn’s post hoc multiple comparisons (C, E, G, and I). Details on number of samples are presented in Table S1.

Overall, these findings demonstrate a clear and novel molecular mechanism of ADM regulation, which restores the disrupted mitochondria-calcium-cytoskeleton-migration axis in 22q11.2DS interneurons.

### ADM’s Rescue of Mitochondrial Deficits and Migration Requires Both PKA and PLC Signaling

To dissect the downstream signaling pathways mediating ADM’s rescue effects, we employed pharmacological inhibitors targeting the two principal arms of ADM signaling: a PKA inhibitor (KT5720; PKAi), a PLC inhibitor (U73122; PLCi), and the competitive receptor antagonist ADM22-52 (Figure 6A). Six experimental conditions were tested: control, 22q11.2DS, 22q11.2DS + ADM, 22q11.2DS + ADM + ADM22-52, 22q11.2DS + ADM + PKAi, and 22q11.2DS + ADM + PLCi.

**Figure 6.**
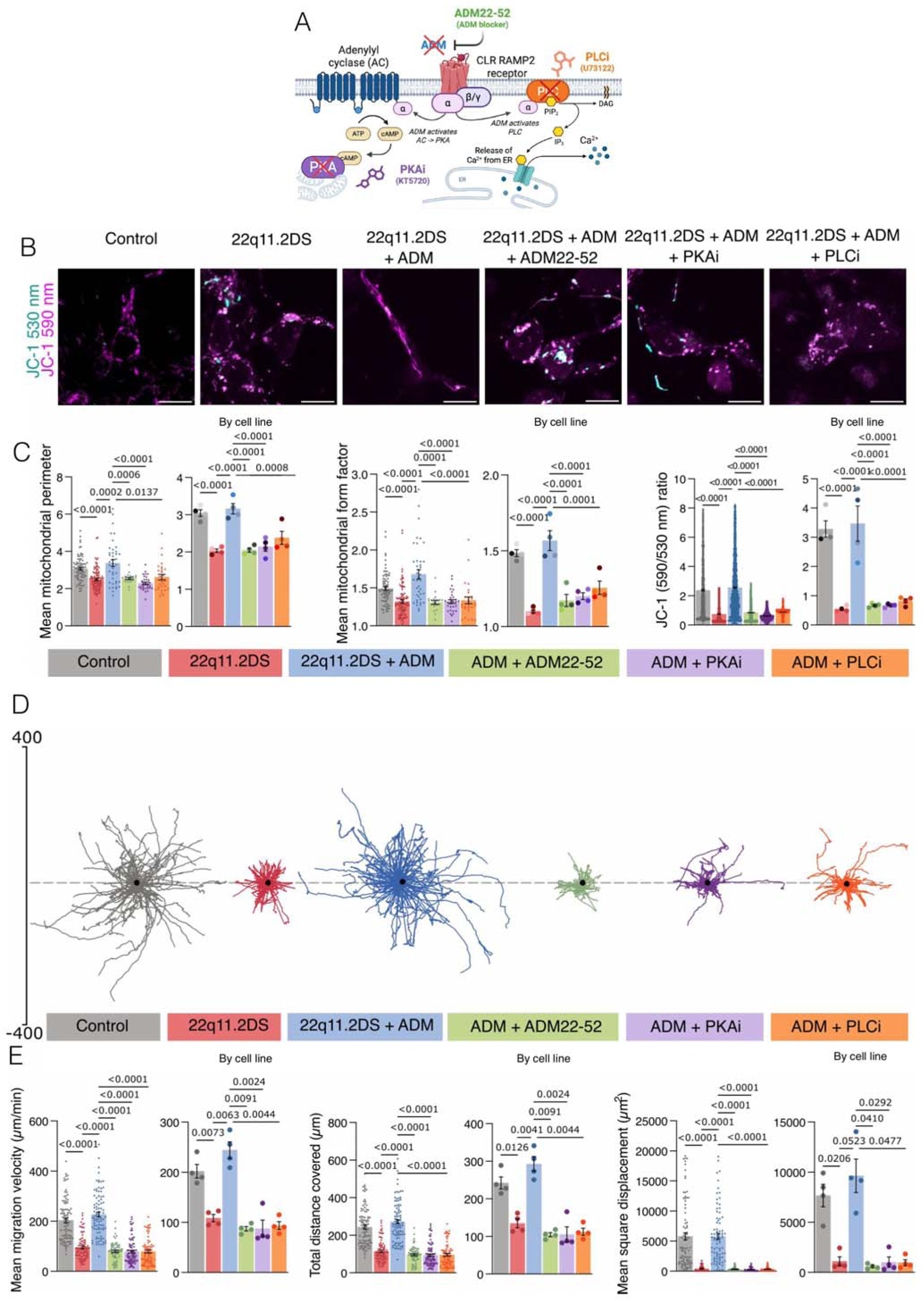
ADM Causes PKA and PLC Pathway Activation to Rescue Mitochondrial Deficits Pharmacological dissection shows that ADM’s rescue of mitochondrial shape, membrane potential, and movement depends on both PKA and PLC signaling, while the receptor antagonist ADM22-52 blocks all rescue effects. **(A)** Diagram of ADM signaling through the CLR/RAMP2 complex: PKA pathway (Gα → AC → cAMP → PKA) and PLC pathway (Gα → PLC → IP₃ → Ca²⁺). Inhibitor sites for PKA inhibitor (KT5720; PKAi), PLC inhibitor (U73122; PLCi), and the competitive receptor antagonist ADM22-52 are marked. **(B)** Typical JC-1 images (530 nm + 590 nm merged) for six conditions: control, 22q11.2DS, 22q11.2DS + ADM, 22q11.2DS + ADM + ADM22-52, 22q11.2DS + ADM + PKAi, and 22q11.2DS + ADM + PLCi. Scale bar, 20 µm. **(C)** Quantification across six conditions: mean mitochondrial perimeter (p < 0.0001, p = 0.0006, p = 0.0002, p = 0.0137, Kruskal-Wallis test with Dunn’s post hoc multiple comparisons), form factor (p < 0.0001, p = 0.0008, Kruskal-Wallis test with Dunn’s post hoc multiple comparisons), perimeter and form factor with inhibitors (p < 0.0001 for multiple comparisons, Kruskal-Wallis test with Dunn’s post hoc multiple comparisons), and JC-1 590/530 nm ratio (p < 0.0001 for multiple comparisons, Kruskal-Wallis test with Dunn’s post hoc multiple comparisons). For mitochondrial morphology, each dot shows one FOV; n = 80 (control), 98 (22q11.2DS), 40 (22q11.2DS + ADM), 20 (22q11.2DS + ADM + ADM22-52), 30 (22q11.2DS + ADM + PKAi), and 28 (22q11.2DS + ADM + PLCi) FOV from 4 iPSC lines per condition. For JC-1 ratio, each dot indicates one mitochondrion; n = 1,669 (control), 2,726 (22q11.2DS), 2,584 (22q11.2DS + ADM), 3,379 (22q11.2DS + ADM + ADM22-52), 2,565 (22q11.2DS + ADM + PKAi), and 1,978 (22q11.2DS + ADM + PLCi) mitochondria from 4 iPSC lines per condition. **(D)** Migration paths: control (gray), 22q11.2DS (red), 22q11.2DS + ADM (blue), 22q11.2DS + ADM + ADM22-52 (green), 22q11.2DS + ADM + PKAi, and 22q11.2DS + ADM + PLCi. Dots mark starting points. **(E)** Migration measurements (pooled and by cell line): speed (µm/min; p < 0.0001, p = 0.0091, p = 0.0024, p = 0.0063, p = 0.0044, p = 0.0073, Kruskal-Wallis test with Dunn’s post hoc multiple comparisons), total distance (p < 0.0001, p = 0.0126, Kruskal-Wallis test with Dunn’s post hoc multiple comparisons), saltation (p < 0.0001, p = 0.0041, p = 0.0044, p = 0.0024, p = 0.0091, Kruskal-Wallis test with Dunn’s post hoc multiple comparisons), and mean square displacement (µm²; p < 0.0001, p = 0.0206, p = 0.0292, p = 0.0410, p = 0.0523, p = 0.0477, Kruskal-Wallis test with Dunn’s post hoc multiple comparisons). Each dot depicts one tracked cell; n = 96 (control), 92 (22q11.2DS), 94 (22q11.2DS + ADM), 66 (22q11.2DS + ADM + ADM22-52), 87 (22q11.2DS + ADM + PKAi), and 76 (22q11.2DS + ADM + PLCi) cells from 4 iPSC lines per condition. Data are shown as mean ± SEM. See also Figure S5.

JC-1 imaging across all six conditions demonstrated that ADM’s rescue of MMP was fully blocked by ADM22-52, PKAi, and PLCi (Figures 6B and 6C; Figure S5A). Similarly, mitochondrial morphology analysis showed that ADM’s rescue of perimeter (p < 0.0001, p = 0.0006, p = 0.0002, p = 0.0137) and form factor (p < 0.0001, p = 0.0008) was abrogated by all three inhibitors, with JC-1 590/530 nm ratios confirming these results (p < 0.0001 for multiple comparisons) (Figure 6C). These experiments demonstrate that the rescue by ADM is mediated by its binding to the RAMP2 receptor, and align with extensive previous literature, including from our group^20^; they also demonstrate that both PKA and PLC pathways are important for this rescue in 22q11.2DS.

In line with the data above, migration analysis across six conditions revealed that ADM’s rescue of interneuron motility was similarly dependent on both PKA and PLC signaling. ADM22-52, PKAi, and PLCi each blocked ADM’s rescue of velocity (µm/min; p < 0.0001, p = 0.0091, p = 0.0024, p = 0.0063, p = 0.0044, p = 0.0073), saltation (p < 0.0001, p = 0.0041, p = 0.0044, p = 0.0024, p = 0.0091), total distance (p < 0.0001, p = 0.0126), and mean square displacement (µm²; p < 0.0001, p = 0.0206, p = 0.0292, p = 0.0410, p = 0.0523, p = 0.0477) (Figures 6D and 6E). These results demonstrate that both the PKA and PLC arms of ADM signaling are required for the full rescue of mitochondrial phenotypes and interneuron migration. Significance was tested with Kruskal-Wallis and Dunn’s post hoc test (C and E). Details on the number of samples are presented in Table S1.

### PLC but Not PKA Mediates ADM’s Rescue of Calcium Transients and Actin Retrograde Flow

Since both PKA and PLC are required for ADM’s rescue of mitochondrial and migration phenotypes based on the above results, we next asked whether these pathways have separable roles in calcium and cytoskeletal regulation. Calcium imaging across six conditions revealed a striking dissociation: PLCi blocked ADM’s rescue of calcium transients, producing significantly reduced peak amplitude and AUC comparable to untreated 22q11.2DS levels (Figure 7A).

**Figure 7.**
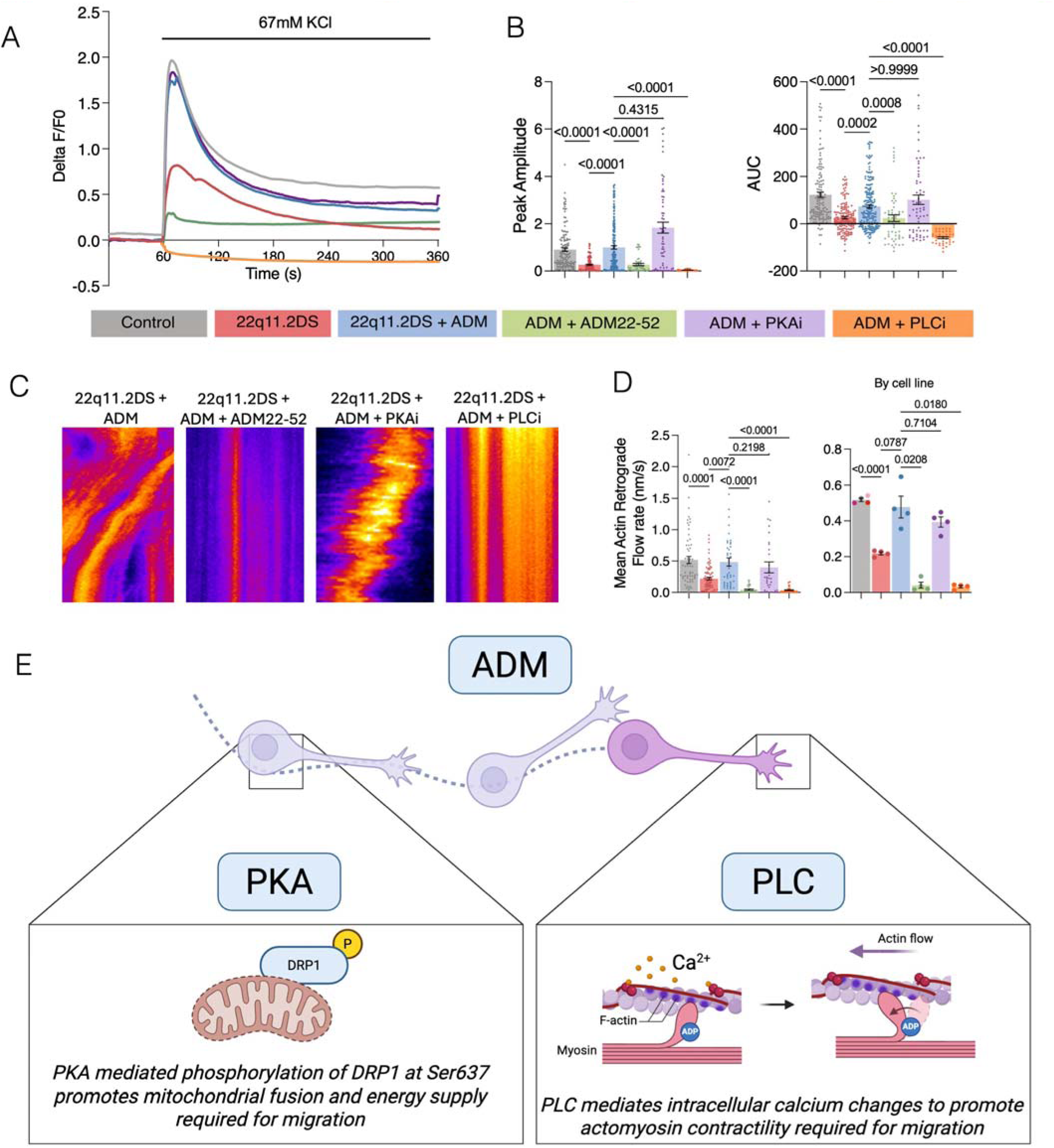
Rescue of Calcium and Cytoskeletal Defects by ADM Are Mediated by the PLC Pathway PLC inhibition, but not PKA inhibition, blocks ADM’s rescue of calcium transients and actin retrograde flow, demonstrating that the PLC pathway mediates ADM’s effects on calcium homeostasis and cytoskeletal dynamics in 22q11.2DS interneurons. **(A)** Calcium transient traces (ΔF/F₀, 67 mM KCl stimulation) across six conditions: control (gray), 22q11.2DS (red), 22q11.2DS + ADM (blue), 22q11.2DS + ADM + ADM22-52, 22q11.2DS + ADM + PKAi, and 22q11.2DS + ADM + PLCi over 360 s. **(B)** Quantification of peak amplitude (p < 0.0001, p = 0.4315, Kruskal-Wallis test with Dunn’s post hoc multiple comparisons) and AUC (p < 0.0001, p > 0.9999, Kruskal-Wallis test with Dunn’s post hoc multiple comparisons). PLCi blocks ADM rescue of calcium (significant reduction); PKAi does not (p = 0.4315, Kruskal-Wallis test with Dunn’s post hoc multiple comparisons). Each dot represents one neuron; n = 139 (control), 152 (22q11.2DS), 212 (22q11.2DS + ADM), 50 (22q11.2DS + ADM + ADM22-52), 66 (22q11.2DS + ADM + PKAi), and 40 (22q11.2DS + ADM + PLCi) neurons from 4 iPSC lines per condition. **(C)** Representative actin retrograde flow kymographs across conditions: 22q11.2DS + ADM, 22q11.2DS + ADM + ADM22-52, 22q11.2DS + ADM + PKAi, and 22q11.2DS + ADM + PLCi. **(D)** Actin dynamics quantification: mean retrograde flow rate (p < 0.0001, p = 0.0087, p = 0.0001, p = 0.0096, Kruskal-Wallis test with Dunn’s post hoc multiple comparisons) and additional parameters (p = 0.0189, p = 0.0787, p = 0.0002, p = 0.0006, Kruskal-Wallis test with Dunn’s post hoc multiple comparisons). Each dot represents one growth cone; n = 60 (control), 83 (22q11.2DS), 46 (22q11.2DS + ADM), 34 (22q11.2DS + ADM + ADM22-52), 38 (22q11.2DS + ADM + PKAi), and 36 (22q11.2DS + ADM + PLCi) growth cones from 4 iPSC lines per condition. **(E)** Summary model. ADM activates two parallel downstream pathways via the CLR/RAMP2 receptor complex: (1) the PKA pathway phosphorylates DRP1 at Ser637, promoting mitochondrial fusion and energy supply for migration; (2) the PLC pathway mediates intracellular calcium changes, promoting actomyosin contractility and actin retrograde flow for migration. Data are represented as mean ± SEM. n values are as indicated per panel from 4 iPSC lines per condition.

Actin retrograde flow analysis mirrored this pattern. PLCi blocked ADM’s restoration of actin dynamics, while PKAi did not (Figures 7C and 7D). Quantification of mean retrograde flow rate confirmed that PLCi significantly impaired ADM’s rescue (p < 0.0001, p = 0.0087, p = 0.0001, p = 0.0096), while additional flow parameters supported this conclusion (p = 0.0189, p = 0.0787, p = 0.0002, p = 0.0006) (Figure 7D). Statistical significance was determined by Kruskal-Wallis test with Dunn’s post hoc multiple comparisons (B and D). Details on number of samples are presented in Table S1.

These findings establish a novel dual-pathway model for ADM’s mechanism of action (Figure 7E). The PKA pathway phosphorylates DRP1 at Ser637, promoting mitochondrial fusion and thereby restoring the bioenergetic capacity required for migration. The PLC pathway mediates intracellular calcium changes, promoting actomyosin contractility and actin retrograde flow, which are essential for growth cone motility and migration. Both arms converge on rescuing interneuron migration through mechanistically distinct routes: PKA provides the energetic substrate via mitochondrial fusion, while PLC provides the cytoskeletal machinery through calcium-dependent actin dynamics.

## DISCUSSION

In this study, we demonstrate that genes associated with neuropsychiatric diseases are preferentially differentially expressed in inhibitory neurons in 22q11.2DS, and, separately, that mitochondrial dysfunction also looks enriched in inhibitory neuronal lineages in 22q11.2DS. Next, we show that human subpallial organoids and forebrain assembloids derived from 22q11.2DS patient iPSCs exhibit a coupled mitochondrial-migratory deficit characterized by mitochondrial fragmentation, reduced oxidative phosphorylation, impaired calcium signaling, disrupted actin dynamics, and deficient interneuron migration. We identify ADM as a potent rescue agent that simultaneously restores mitochondrial morphology, membrane potential, respiration, calcium transients, actin retrograde flow, and migration. Mechanistic dissection reveals that ADM engages two parallel signaling cascades, PKA and PLC, with separable downstream effects: PKA drives DRP1 Ser637 phosphorylation and mitochondrial fusion, while PLC restores calcium homeostasis and actin dynamics. These findings, validated in primary human cortical tissue carrying the 22q11.2 deletion, establish a mitochondrial cytoskeletal axis as a convergent disease mechanism and identify ADM and its downstream pathways as candidates for therapeutic discovery for neuropsychiatric and global mitochondrial pathology in 22q11.2DS.

Our transcriptomic and functional data converge on mitochondrial dysfunction as a central feature of interneuron pathology in 22q11.2DS, thus strengthening the existing literature about the mitochondrial dysfunction in 22q11.2DS^10,15–17,30^. The resulting imbalance in DRP1 phosphorylation, elevating Ser616 (pro-fission) and reducing Ser637 (pro-fusion)^31^, provides a mechanistic link between haploinsufficiency of mitochondrial genes and the fragmented mitochondrial phenotype. DRP1 has emerged as a critical regulator in multiple neurodevelopmental contexts, and our findings position it as a key therapeutic node in 22q11.2DS^31–35^.

Separately, the calcium signaling defect we observe in interneurons has also been described in excitatory neurons from 22q11.2DS^12^, suggesting a more global effect of the deletion on multiple types on neurons.

The migration deficits we identify in both hFAs and primary hCT carry significant implications for cortical circuit formation in 22q11.2DS. Interneurons must traverse long distances from their subpallial origins to reach appropriate cortical positions, a process that demands sustained bioenergetic output, precise calcium signaling, and dynamic cytoskeletal remodeling^36–40^. Impaired migration could result in reduced cortical interneuron density, aberrant laminar positioning, and ultimately disruption of the excitation-inhibition balance, a feature increasingly recognized as a core feature of schizophrenia and other neuropsychiatric disorders^19,20,39,41–44^. Our findings align with prior observations in mouse models of 22q11.2 deletion, which demonstrated interneuron positioning defects^30,45,46^ and indications of mitochondrial dysfunction and impaired interneuron migration in a recently published forebrain assembloid model containing PVALB interneurons^42^. The concordance between organoid and hCT phenotypes underscores the fidelity of the organoid model and highlights the value of human-specific experimental systems for studying neurodevelopmental disorders.

ADM’s dual engagement of PKA and PLC signaling provides a uniquely comprehensive rescue that addresses both the energetic and cytoskeletal dimensions of the migratory deficit. The PKA arm, by phosphorylating DRP1 at Ser637, promotes mitochondrial fusion and restores membrane potential, effectively replenishing the bioenergetic supply required for sustained migration. The PLC arm, operating in parallel with the PKA arm (and as shown by our pharmacological dissection, also contributing to mitochondrial rescue), restores intracellular calcium transients and normalizes actin retrograde flow, thereby re-engaging the cytoskeletal machinery that powers growth cone motility. The requirement of both pathways for full migration rescue reveals that interneuron migration in 22q11.2DS is accompanied by concurrent mitochondrial and cytoskeletal deficits, each of which must be independently corrected. ADM has an established clinical safety profile in the Crohn’s disease literature^23^, and its ability to engage both arms simultaneously makes it an appealing candidate for translational development. Key challenges for clinical translation include optimizing dosing regimens, defining treatment windows during development, and identifying routes of delivery to the central nervous system^47,48^.

There are a few important limitations of this study that should be considered. Although we examined four iPSC lines per genotype to obtain robust, convergent phenotypes across patients, more iPSC lines will be needed in future work to capture the full spectrum of 22q11.2DS, which exhibits highly heterogeneous disease manifestations. Furthermore, although organoid models recapitulate key aspects of interneuron development and migration, they currently lack vascularization, immune interactions, and long-distance circuit connectivity. *Ex vivo* human developing cortical tissue experiments, though providing critical human validation, were limited by tissue availability and represent a single developmental time point. The ADM concentration and treatment window used in this study have not been optimized, and dose-response relationships, potential off-target effects, and long-term consequences of ADM treatment remain to be characterized.

In conclusion, this study confirms that disruption of the mitochondrial-cytoskeletal axis is a convergent disease mechanism underlying interneuron dysfunction in 22q11.2DS and identifies ADM as a potential molecule for correction. By showing consistent phenotypes and rescue effects in both iPSC-derived organoids and primary human cortical tissue, we lay the groundwork for developing mitochondrial-targeted therapies for neuropsychiatric issues associated with 22q11.2DS and, more broadly, for addressing the challenges of multi-organ mitochondrial defects in the condition.

Future studies should investigate ADM’s effects *in vivo*, optimize therapeutic parameters, and explore whether this mitochondrial migratory axis is disrupted in other neurodevelopmental conditions with mitochondrial involvement, potentially broadening the therapeutic relevance of our findings.

## RESOURCE AVAILABILITY

### Lead Contact

Further information and requests for resources and reagents should be directed to and will be fulfilled by the lead contacts, Anca Pasca and Dhriti Nagar

### Materials Availability

This study did not generate new, unique reagents.

### Data and Code Availability

The Walsh dataset is publicly available under accession GSE250482 (Walsh et al., Neuron, 2026). The CellAtria framework (v1.0.0) and the CellExpress pipeline are publicly available at https://github.com/AstraZeneca/cellatria under an open-source license. Pipeline configuration files (JSON) capturing all runtime parameters for each module have been deposited alongside the processed data. Any additional information required to reanalyze the data reported in this paper is available from the lead contact upon request.

## Supporting information

Key Resource Table

## ACKNOWLEDGMENTS

We thank the members of the Brain Therapeutics Laboratory at Stanford University for helpful discussions and critical reading of the manuscript. We are grateful to Dr. Valentin Cracan for valuable advice regarding Seahorse extracellular flux analyses. We thank the staff of the Stanford Shared FACS Facility for their assistance with flow cytometry and cell sorting, and the Weinacht Laboratory for access, training and experimental guidance with the Seahorse XF analyzer. We thank the donors and the Department of Obstetrics and Gynecology at Stanford Hospital for providing access to human cortical tissue. This work was supported by a Maternal & Child Health Research Institute (MCHRI) Postdoctoral Fellowship to D.N., Uytengsu-Hamilton 22q11 Neuropsychiatry Research Award (MCHRI) to A.M.P., McCormick & Gabilan Faculty Award, Stanford to A.M.P., Dunlevie Maternal-Fetal Medicine Center for Discovery, Innovation, and Clinical Impact, Stanford to A.M.P.

## AUTHOR CONTRIBUTIONS

D.N. and A.M.P. conceptualized and designed the study. D.N. performed most experiments, including iPSC culture and differentiation, generation of cortical organoids and assembloids, ADM treatment studies, calcium and mitochondrial imaging, and biochemical validation. D.N. carried out formal analysis, generated the figures, and wrote the original draft of the manuscript. J.C. contributed to the investigation and methodology, including Western blots for DRP1 phosphorylation status. L.L. contributed to the methodology and analysis of flow cytometry experiments. M.S.H., S.P., B.A., A.N., M.A., and S.H. contributed to the investigation, including iPSC maintenance, organoid generation, quantitative PCRs, and image quantification. Z.H. performed computational analysis of sequencing data and contributed to data curation under the supervision of Z.G. Z.G. supervised computational analyses, contributed to methodology, and provided resources. W.W. contributed investigation and methodology related to Seahorse assays under the supervision of K.G.W. K.G.W. provided resources, contributed to methodology, and reviewed the manuscript. A.M.P. supervised the study, acquired funding, provided resources, contributed to project administration, and wrote and edited the manuscript with D.N. All authors reviewed and approved the final manuscript.

## METHODS

### scRNA-seq Data Reanalysis

The scRNA-seq dataset was obtained from the publicly available dataset on brain assembloids harboring a 22q11.2 deletion from Walsh et al. (2026, Neuron; GEO: GSE250482). This dataset comprises scRNA-seq from hiPSC-derived forebrain assembloids generated by fusing dorsally patterned cortical organoids with ventrally patterned ganglionic eminence organoids, profiled at day 60 and beyond. Dataset was retrieved and processed using CellAtria (v1.0.0; Nouri et al., npj Artificial Intelligence, 2026), an agentic AI framework that automates GEO data retrieval, file standardization to 10X Genomics conventions, and pipeline execution via a graph-based multi-actor architecture with 32 callable tools.

Raw count matrices were processed through the CellExpress (v1.0.0) Scanpy-based pipeline. Cells were retained if they contained ≥750 UMI counts and ≥250 detected genes, with mitochondrial transcript fraction <15%; doublets were removed using Scrublet (score threshold 0.25). Counts were library-size normalized (target sum 10,000) and log1p-transformed, and the top 2,000 highly variable genes were selected for downstream analysis. PCA was performed (30 components), followed by Harmony batch correction on sample identity. UMAP embeddings were computed on the corrected space, and Leiden clustering (resolution 0.6) was applied to a k = 15 nearest-neighbor graphs. Cell type annotation was performed using SCimilarity (v1.1) as a tissue-agnostic classifier and CellTypist (v1.6.3) with the Developing_Human_Brain model, with cluster-level assignments by majority vote. Cluster marker genes were identified by Wilcoxon rank-sum test (adjusted p ≤ 0.05, log_₂_FC ≥ 0.25, detection in ≥10% of cluster cells).

### Human iPSC Lines

Four control iPSC lines (2242-1, 1205-4, 1208-2, 8119-1) and four 22q11.2DS patient-derived iPSC lines (6303-5, 7958-3, 1804-5, 9050-3) were used in this study. These iPSC lines have been previously validated with confirmed deletion and normal karyotype (Khan et al., 2020)^20^ and were obtained from the laboratory of Sergiu Pasca. All iPSC work was performed under institutional review board (IRB) approval at Stanford University (Protocol SCRO-796). The iPSCs were cultured on recombinant human Vitronectin (rh-VTN; Thermo Fisher Scientific, Cat#A14700) and maintained in Essential 8 medium (Thermo Fisher Scientific, Cat#A1517001) with EDTA-based gentle passaging of colonies. The cells were grown to 80% confluency before generating organoids.

### Human Subpallial Organoid (hSO) Generation

hSOs were generated using a feeder-free ventral forebrain patterning protocol (Puno et al., 2026). Briefly, iPSC colonies were dissociated into single cells using Accutase (Innovate Cell Technologies, Cat#AT-104) and counted. The dissociated cells were aggregated into embryoid bodies (10,000 cells per EB) in AggreWell plate-800 (STEMCELL Technologies, Cat#34815) in Essential 8 medium (Thermo Fisher Scientific, Cat#A1517001) supplemented with CEPT cocktail (Selleck Chemicals, Cat#S1049). After successful aggregation, the embryoid bodies were transferred to Essential 6 medium (Thermo Fisher Scientific, Cat#A1516401) containing Dorsomorphin (DM at 5 μM final concentration; Sigma-Aldrich, Cat#P5499), SB431542 (SB at 10 μM final concentration; Tocris, Cat#1614), and XAV-939 (XAV at 2.5 µM final concentration; Tocris, Cat#3748) for neural induction with dual SMAD inhibition. On day 6, the media was changed to neural media (Neurobasal A (Thermo Fisher Scientific, Cat#10888022) with B-27 supplement without Vitamin A (Thermo Fisher Scientific, Cat#12587010), 1X GlutaMax (Thermo Fisher Scientific, Cat#35050061), and 1X PenStrep (Thermo Fisher Scientific, Cat#15140122)). Neural progenitor proliferation was achieved using EGF (20 ng/ml; R&D Systems, Cat#236-EG), FGFb (20 ng/ml; R&D Systems, Cat#233-FB), and XAV-939 (2.5 μM; Tocris, Cat#3748) until day 23, with daily medium changes during the first 10 days and every other day thereafter. Ventral patterning was maintained through supplementation of organoids with SAG (Thermo Fisher Scientific, Cat#566660) from day 12 to day 23 in culture. Starting on day 25, media were supplemented with BDNF (20 ng/ml; Peprotech, Cat#450-02) and NT3 (20 ng/ml; Peprotech, Cat#450-03) to promote neuronal maturation, and media were changed every other day until day 43. Organoids were maintained in neural media, with media changes every 3-4 days, until day 100.

### Bulk RNA Sequencing and Analysis

Day 100 hSOs generated from Control and 22q11.2DS lines were collected in DNA/RNA Shield solution (Zymo Research). Bulk RNA sequencing was performed by Plasmidsaurus using a 3’ end counting approach. Total RNA was extracted from human cell samples preserved in Zymo DNA/RNA Shield using a bead-based extraction method. mRNA transcripts were converted to cDNA via reverse transcription using a poly(dT)VN primer, followed by second-strand synthesis and tagmentation. Sequencing libraries were indexed using unique dual indices (UDIs) and amplified. Sequencing was conducted on the Illumina platform with single-end reads of approximately 90 bp, targeting the 3’ end of transcripts (within the final ∼400 nt). Unique molecular identifiers (UMIs) were utilized during library preparation to facilitate accurate computational deduplication.

### Bioinformatics Pipeline (Plasmidsaurus)

Quality of the fastq files was assessed using FastQC v0.12.1. Reads were then quality filtered using fastp v0.24.0 with poly-X tail trimming, 3’ quality-based tail trimming, a minimum Phred quality score of 15, and a minimum length requirement of 50 bp. Filtered reads were aligned to the Human Reference Genome (GRCh38) using STAR v2.7.11 with non-canonical splice junction removal and output of unmapped reads, followed by coordinate sorting using samtools v1.22.1. PCR and optical duplicates were removed using UMI-based deduplication with UMIcollapse v1.1.0. Alignment quality metrics, strand specificity, and read distribution across genomic features were assessed using RSeQC v5.0.4 and Qualimap v2.3, and the results were aggregated into a comprehensive quality control report using MultiQC v1.32. Gene-level expression quantification was performed using featureCounts (subread package v2.1.1) with strand-specific counting, multi-mapping read fractional assignment, exons, and three prime UTR as the feature identifiers, and grouped by gene_id. Final gene counts were annotated with gene biotype and other metadata extracted from the reference GTF file. Sample-sample correlations for the sample-sample heatmap and PCA were calculated on normalized counts (TMM, trimmed mean of M-values) using Pearson correlation. Differential expression was performed with edgeR v4.0.16 using standard practice, including filtering for lowly expressed genes with edgeR::filterByExpr using default values. Functional enrichment, when available, is performed using gene set enrichment analysis with gseapy v0.12 using the MSigDB Hallmark gene set.

### Neuropsychiatric Risk Gene Set Overlap Analysis

DEGs from 22q11.2DS versus control hSOs were compared with risk gene sets from ADHDgene (ADHD), PGC (MDD and bipolar disorder), SZDB (schizophrenia), and SFARI (autism) as available on DisGeNet. Overlap significance was determined by Fisher’s exact test.

### Calcium Imaging

Dissociated hSOs were plated on Poly L-Ornithine (Sigma) coated coverslip bottom 96-well plates at a density of 20,000 cells per well. After 48 hours of outgrowth in neural media, cells were incubated in CalBryte-520 AM (AAT BioQuest, Cat#20650; 5 µM) for 30 minutes at 37 °C. The cells were then washed and incubated in base Tyrode solution. The base Tyrode’s solution was prepared containing 129 mM NaCl, 2 mM CaCl_₂_, 1 mM MgCl_₂_, 30 mM Glucose, and 25 mM HEPES. To establish baseline conditions, KCl was added to a concentration of 5 mM. For experiments conducted in 96-well plates, each well contained 100 µl of the baseline solution. Chemical stimulation was achieved by adding 100 µl of a 2X high-potassium Tyrode’s solution (129 mM KCl) to reach a final working concentration of 67 mM KCl. Each round of calcium imaging included a 60-second baseline, stimulation with 2X high-potassium Tyrode’s solution at the 60-second mark, and imaging for 300 seconds thereafter. Fluorescence intensity changes were calculated as ΔF/F_₀_, and peak amplitude and AUC were quantified.

### Flow cytometry analysis of organoid composition

hSOs were dissociated into a single suspension, filtered through a 70µm strainer, and centrifuged to obtain a cell pellet. The cells were incubated in Zombie Violet 405 fixable viability dye (Biolegend, Cat#423113; 1:1000 in DPBS (Thermo Fisher Scientific, Cat#14190144) without Magnesium and Calcium; 0.2% BSA) for 15 minutes. The cells were fixed and permeabilized using the BD Cytofix/Cytoperm kit as per the manufacturer’s instructions. Permeabilized cells were blocked for 30 minutes in blocking buffer (Cytoperm buffer containing 5% BSA, 5% Normal Donkey serum, and 0.01% Sodium Azide). Intracellular staining of the cells was performed in the blocking buffer containing primary antibodies to stain progenitors (goat-anti-human SOX2; 1:500) and neurons (mouse-anti-beta-III-tubulin (TUJ1; Biolegend, Cat#801202); 1:500) for 30 minutes at 4 °C. Cells were washed thrice for 5 minutes each in the Cytoperm buffer. Washed cells were incubated in blocking buffer containing secondary antibodies, donkey-anti-goat AlexaFluor488 (Invitrogen, Cat#A-11055), and donkey-anti-mouse AlexaFluor647 (Invitrogen, Cat#A-31571). 200,000 live singlet cells per condition per replicate were analyzed using the Quanteon or Penteon Bioanalyzer (Agilent Technologies).

### Human Cortical Organoid (hCO) Generation

hCOs were generated using a dorsal forebrain patterning protocol (Puno et al., 2026). Briefly, the hSO generation protocol was followed till day 5. At day 6, neural progenitor proliferation was achieved using EGF (20 ng/ml; R&D Systems, Cat#236-EG) and FGFb (20 ng/ml; R&D Systems, Cat#233-FB) until day 23, with daily medium changes during the first 10 days and every other day thereafter. Starting on day 25, media were supplemented with BDNF (20 ng/ml; Peprotech, Cat#450-02) and NT3 (20 ng/ml; Peprotech, Cat#450-03) to promote neuronal maturation, and media were changed every other day until day 43. Organoids were maintained in neural media, with media changes every 3 to 4 days, until day 100.

### Human Forebrain Assembloid (hFA) Generation

hFAs were generated by placing one hSO (transduced with Dlx1/2i::EGFP lentivirus at day 50) and one hCO in a low attachment well at day 60 of differentiation. Fusion was allowed to proceed for 15 days. Live imaging of interneuron migration was performed from day 75 onward.

### Lentiviral Production and Transduction

The Dlx1/2i::EGFP reporter construct was packaged into lentiviral particles using pVSV-G, pMDL, and pRSV in HEK293T cells (ATCC, Cat#CRL-3216; RRID: CVCL_0063). Viral supernatant was collected at 48 hours post-transfection and concentrated using a Lenti-X concentrator (Lonza). hSOs or cortical slices were transduced for 10 days and 4 days, respectively.

### FACS of Dlx1/2i::eGFP+ Interneurons

hSOs at day 100 were dissociated into single-cell suspensions using Accutase (Innovate Cell Technologies, Cat#AT-104) at 37 °C for 40 minutes, followed by gentle trituration. Dissociated cells were rinsed twice with media containing CEPT (Selleck Chemicals, Cat#S1049). Cells were filtered through a 70 µm strainer to remove debris, resuspended in sorting buffer containing NucBlue viability stain, and sorted for EGFP(+) and NucBlue(-) cells using a BD Fusion FACS instrument (100 µm microfluidic chip).

### Quantitative RT-PCR

RNA was extracted using the RNeasy Plus kit (Qiagen, Cat#74136) and reverse-transcribed using the SuperScript IV cDNA synthesis kit (Thermo Fisher Scientific, Cat#18091050). qPCR was performed on a BIORAD CFX96 instrument using SYBR Green master mix (Vazyme). Primers are listed in the key resources table. Expression was normalized to GAPDH using the ΔCt method.

### Seahorse XF Mito Stress Test

Cellular respiration was measured using a Seahorse XFe96 Analyzer (Agilent). hSOs were dissociated, and 50,000 cells were seeded per well and maintained in neural media. On the day of the assay, culture media were replaced with 180 ul of Seahorse Media (DMEM) supplemented with 1 mM Glutamine, 1 mM Pyruvate, and 2.5 mM Glucose. Cells were incubated in a non-CO[ incubator at 37°C for 60 minutes prior to the start of the assay. Baseline oxygen consumption rates (OCR) were measured for 30 minutes (5 cycles) followed by sequential injections of metabolic inhibitors to assess mitochondrial function. The Mito Stress Test was performed using the following final concentrations: 5 µM Oligomycin (Sigma-Aldrich, Cat#75351; Port A) to inhibit ATP synthase, 4 µM FCCP (Sigma-Aldrich, Cat#C2920; Port B) to uncouple oxidative phosphorylation, and 3 µM Rotenone (Cayman Chemicals, Cat#13995; Port C) to inhibit Complex I. Each drug phase consisted of three 6-minute cycles, including a 3-minute mixing period and a 3-minute measurement period, for a total duration of 18 minutes (Oligomycin), 18 minutes (FCCP), and 18 minutes (Rotenone). After the run was complete, the BCA assay (Pierce™ BCA Protein Assay Kit; Thermo Fisher Scientific, Cat#23225) was performed and used to normalize the OCR measurements in the Seahorse Analytics web app. Basal respiration, maximal respiration, spare respiratory capacity, non-mitochondrial respiration, ATP production, and proton leak were calculated according to standard protocols.

### Confocal Imaging

Images were acquired on an LSM980 confocal microscope (Zeiss) using a 40X water-immersion objective with a 1.2 NA for all the imaging experiments. Pharmacological treatments were performed 24 hours prior to the imaging assays. Media was replaced with fresh neural media with the drugs for the duration of imaging. Treatments and imaging for different conditions and drug treatments were staggered to account for imaging time and to maintain uniform treatment durations.

### Mitochondrial Morphology Analysis

Cells were stained with pkMitoDeepRed (Spirochrome, Cat#SC055) at 200 nM for 15 minutes at 37 °C with 5% CO[. Cells were washed with fresh medium and incubated at 37 °C and 5% CO[ for 30 minutes prior to imaging. Confocal images were acquired at 1.4 Airy Units. Images were analyzed using Mitochondria Analyzer^49^, a plugin in FIJI. Mitochondria were segmented by reiterative thresholding in batch mode, using the same settings across the entire dataset, blind to genotype or treatment, and morphometric parameters were calculated: perimeter and form factor (perimeter²/[4π × area]). A value of 1 indicates a perfect circle.

### Live Interneuron Migration Imaging and Tracking

hFAs or cortical slices were placed in an environmental chamber (37°C, 5% CO[; OkoLab) and imaged with a.\ Zeiss LSM980 confocal microscope. Dlx1/2i::EGFP labeled interneurons on the surface of the hFAs were imaged every 20 minutes for 24 hours as a 200 µm thick optical section with 4 µm interval using a 10X objective. Maximum-intensity projections along the Z-stack were generated using Zen Blue software. The projection time series was stabilized using the MultiStackReg plugin in FIJI. Cell tracking was performed using the Manual Tracking plugin in FIJI. Migration velocity (µm/min), total distance (µm), and mean square displacement (mm² or µm²) were quantified using the Ibidi chemotaxis tool. Tracks were plotted from a common origin for visualization by subtracting the starting position coordinates from the subsequent coordinates.

### Human Cortical Tissue experiments

Gestational age 19 weeks 4 days (19w4d), human cortical tissue was obtained with CVS confirmation of 22q11.2 deletion. Also,gestational age 19 weeks 2 days (19w2d), human cortical tissue without the 22q11.2 deletion was obtained for control experiments. Tissues were procured under the SCRO-796 approval. Cortical areas were embedded in low-melting agarose (Invitrogen, Cat#16520-100) and uniformly sliced into 2mm sections. These slices were cultured ex vivo for 6 days on cell culture inserts in 6-well ultra-low attachment plates (Corning, Cat#430293) in cortical media. The human cortical (hCT) medium was prepared using Basal Medium Eagle (BME; Gibco, Cat#21010046) as the primary nutrient base. The medium was supplemented with 5% (v/v) Fetal Bovine Serum (FBS; Gibco, Cat#16000044), 25% (v/v) Hanks’ Balanced Salt Solution (HBSS; Gibco, Cat#14025092), 1% (v/v) Penicillin-Streptomycin (Thermo Fisher Scientific, Cat#15140122), 1% (v/v) Antibiotic-Antimycotic, 1% (v/v) GlutaMAX (Thermo Fisher Scientific, Cat#35050061), and 1% (v/v) N2 supplement (Gibco, Cat#17502048). D-glucose (Sigma-Aldrich, Cat# G7021) was added to the formulation freshly at a final concentration of 0.66% (w/v). Ex vivo human cortical slices were transduced with Dlx1/2i::EGFP lentivirus and maintained in hCT medium for 6 days, with daily media changes, for live-tracking experiments. One of the sections was dissociated using Accutase (Innovate Cell Technologies, Cat#AT-104) for 45 minutes, and a single-cell suspension was created by gentle trituration. The dissociated cells were plated on Poly-L-Ornithine and laminin-coated 8-well chamber slides with a coverslip bottom (Ibidi) for mitochondrial imaging assays.

### Adrenomedullin Treatment

Adrenomedullin peptide (Anaspec, Cat#AS-60447) was obtained from Anaspec, reconstituted in ultra-filtered water, and added to cultures at a concentration of 0.5 μM for 24 hours prior to each assay.

### JC-1 Mitochondrial Membrane Potential Assay

Cells were loaded with JC-1 (Thermo Fisher Scientific, Cat#T3168) to quantify mitochondrial membrane potential (MMP) at 10 ug/mL for 30 minutes at 37 °C and 5% CO_₂_. Fluorescence was captured at 530 nm (monomers, green, indicating low MMP) and 590 nm (J-aggregates, red, indicating high MMP). The 590/530 nm ratio was calculated as a measure of MMP. Images were analyzed using the ratiometric intensity analysis tool in the Mitochondria Analyzer plugin in FIJI.

### Actin Retrograde Flow Analysis

Dissociated hSOs plated on Poly-L-Ornithine-coated coverslip bottom 8-well chamber slides were incubated with FastAct-SPY555 (Spirochrome, Cat#SC205) at a 1:2000 dilution for 2 hours at 37°C and 5% CO_₂_ to stain F-actin. Time-lapse imaging was performed at 1 fps for 140 seconds. Kymographs were generated using the KymoResliceWide plugin in FIJI, targeting the growth cone end of migrating neurons. The F-actin retrograde flow rates were calculated from the output kymographs using MATLAB code developed by the Odde lab (2008).

### Pharmacological Inhibitors

PKA inhibitor (PKAi) KT5720 (Tocris Bioscience, Cat#1288) was used at a 0.5 μM concentration. PLC inhibitor (PLCi) U73122 (Tocris Bioscience, Cat#1268) was used at a 0.1 μM concentration. ADM22-52 (Cayman Chemicals, Cat#24892; AM1 receptor competitive antagonist) was used at a 0.5 μM concentration. Inhibitors were added alongside the ADM addition.

### Western Blotting

Whole hSOs were lysed on ice with RIPA buffer supplemented with protease and phosphatase inhibitors (Santacruz, Cat#sc-24948A). The cell lysates were incubated at 4°C for 1 hour and then centrifuged at 14,000 x g for 15 minutes. After centrifugation, the supernatant was collected, and the protein concentration was measured using the BCA Protein Assay Kit (Thermo Fisher Scientific, Cat#23225). The lysate was denatured in Bolt™ LDS Sample Buffer (Invitrogen, Cat#B0007) at 95°C for 5 minutes. Samples with 7 μg input per lane were loaded onto Bolt™ 4-12% Bis-Tris Protein Gels (Invitrogen, Cat#NW04120BOX) and transferred to a PVDF membrane using iBlot™ 2 Transfer Stacks (Invitrogen, Cat#IB24002) and an iBlot 2 (P3) (Thermo Fisher Scientific, Cat#IB21001). After blocking with 5% skim milk (BD, Cat#232100) in TBST (Boston BioProducts, Cat#BM-301X) containing 0.1% Tween 20 (Sigma-Aldrich, Cat#P1379) for 1 hour, the samples were incubated in blocking solution (TBST with 5% BSA (Gendepot, Cat#A0100-005)) containing the primary antibodies at 4°C listed in the resources table. Secondary antibodies conjugated to HRP were used for chemiluminescent detection with SuperSignal™ West Femto Maximum Sensitivity Substrate (Thermo Fisher Scientific, Cat#34095). The blots were imaged with the iBright 1500 imager (Thermo Fisher Scientific, Cat#A44114), and densitometry was performed in FIJI, normalized to total DRP1 and β-actin.

### Quantification and Statistical Analysis

All statistical analyses were performed using GraphPad Prism 10 (GraphPad Software; RRID:SCR_002798). To assess the suitability of the statistical methods, the raw data distribution was evaluated for normality. Two-group comparisons (control vs. 22q11.2DS) were performed using Welch’s t-test (Gaussian data) and Mann-Whitney test (non-Gaussian data). Comparisons among three or more groups were performed using the one-way ANOVA (Gaussian data) and Kruskal-Wallis test with Dunn’s post hoc multiple-comparison correction (non-Gaussian data). Data are represented as mean ± SEM. A significance threshold of p < 0.05 was used. Exact p-values are reported in the figure legends. For each assay, biological replicates represent independent iPSC lines or independent cortical samples, and technical replicates represent individual organoids, cells, or tracks as indicated in the figure legends and Supplementary Table S1.

---

**Figure S1.**
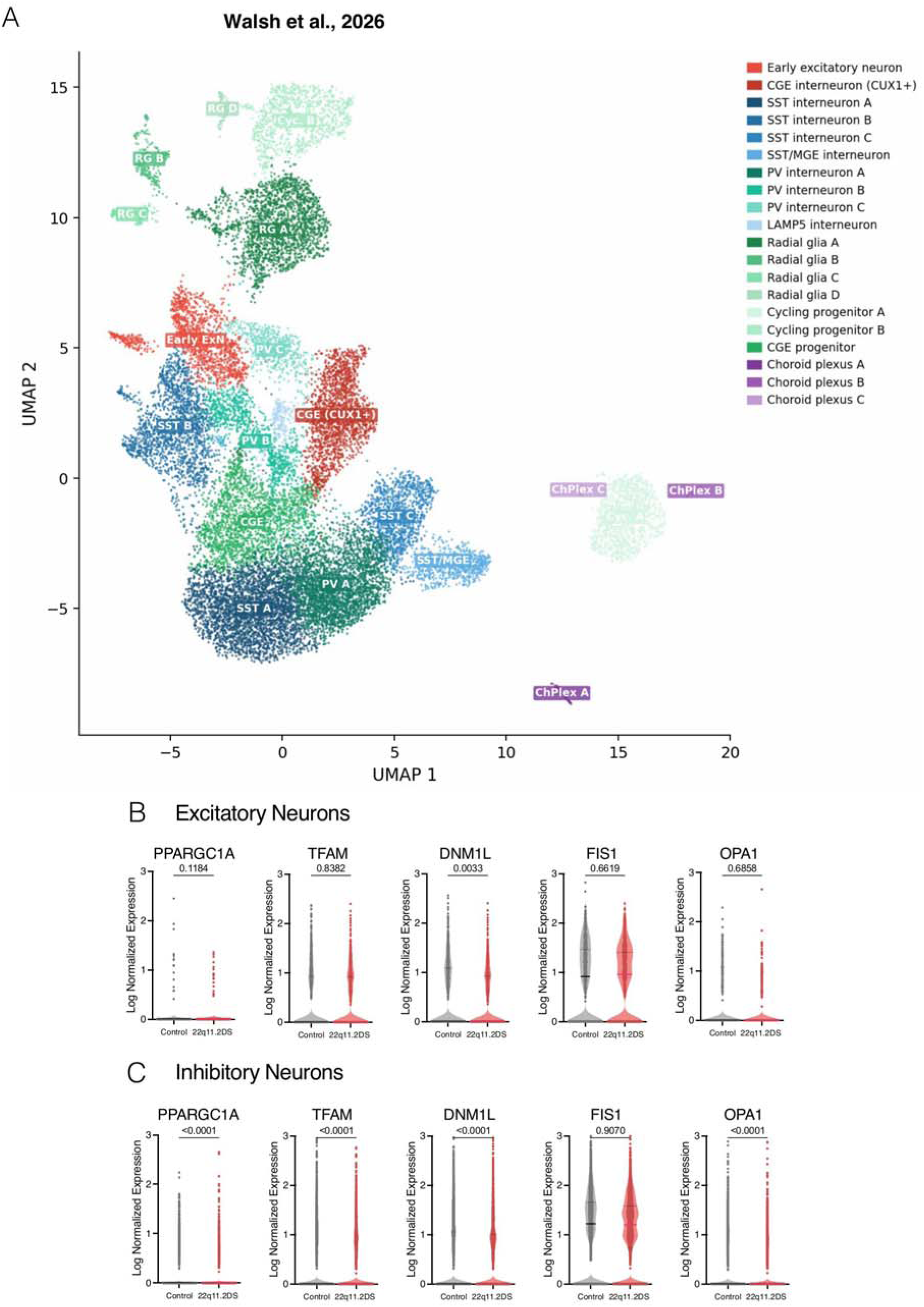
Analysis of Publicly Available 22q11.2DS Brain Assembloids scRNA-seq Dataset Corroborates Enrichment of Mitochondrial Deficits in Inhibitory Neurons, Related to. Figure 1 **Reanalysis of an independently published single-cell RNA-seq (scRNA-seq) dataset from 22q11.2DS brain assembloids reveals cell-type-specific dysregulation of mitochondrial dynamics and biogenesis genes, with preferential vulnerability in inhibitory neurons. (A)** UMAP of scRNA-seq data from Walsh et al., 2026. Violin plots of mitochondrial dynamics and biogenesis gene expression in control (Ctrl, gray) versus 22q11.2DS (22q, red) cells, stratified by cell type: **(B)** Excitatory neurons and **(C)** Inhibitory neurons. PGC1A (PPARGC1A): excitatory (p = 0.118, Wilcoxon rank-sum test), inhibitory (p = 5.6e-08, Wilcoxon rank-sum test). TFAM: excitatory (p = 0.838, Wilcoxon rank-sum test), inhibitory (p = 3.2e-10, Wilcoxon rank-sum test). DNM1L (DRP1): excitatory (p = 0.239, Wilcoxon rank-sum test), inhibitory (p = 1.1e-10, Wilcoxon rank-sum test). FIS1: excitatory (p = 0.662, Wilcoxon rank-sum test), inhibitory (p = 0.907, Wilcoxon rank-sum test). OPA1: excitatory (p = 0.686, Wilcoxon rank-sum test), inhibitory (p = 1.5e-06, Wilcoxon rank-sum test). Y-axes represent log-normalized expression.

**Figure S2.**
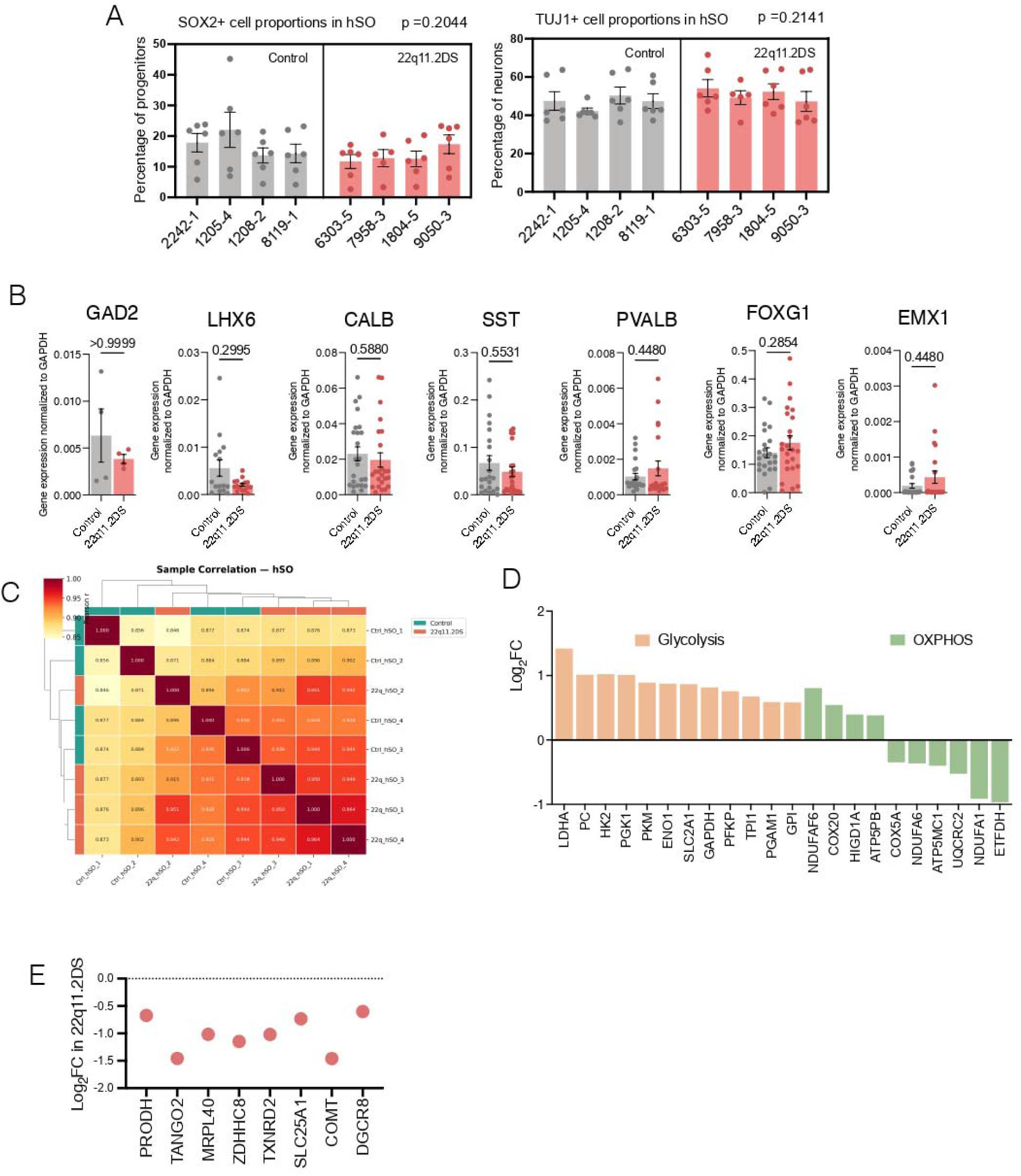
Characterization of 22q11.2DS Human Cellular Models, Related to. Figure 1 **Extended characterization demonstrates preserved progenitor and neuronal proportions across iPSC lines, validates the hemizygous deletion of 22q11.2 locus genes, confirms comparable expression of interneuron subtype markers, and details mitochondrial functional category dysregulation. (A)** Left: quantification of SOX2+ progenitor proportions in control and 22q11.2DS hSOs. Individual iPSC lines are shown on the x-axis (control: 2242-1, 1205-4, 1208-2, 8119-1; 22q11.2DS: 6303-5, 7958-3, 1804-5, 9050-3). p = 0.2044, Welch’s t-test. Right: quantification of TUJ1+ neuron proportions. p = 0.2141, Welch’s t-test. Each dot represents one organoid; n = 12 (control) and 12 (22q11.2DS) organoids from 4 iPSC lines. **(B)** Quantitative reverse-transcription PCR (qRT-PCR) of interneuron subtype markers normalized to GAPDH: GAD2 (p > 0.9999, Welch’s t-test), LHX6 (p = 0.2995, Welch’s t-test), CALB (p = 0.5880, Welch’s t-test), SST (p = 0.5531, Welch’s t-test), PVALB (p = 0.4480, Welch’s t-test), FOXG1 (p = 0.2854, Welch’s t-test), EMX1 (p = 0.4480, Welch’s t-test). Each dot represents one biological replicate: n = 4 iPSC lines per genotype. **(C)** Sample correlation heatmap (Pearson) for bulk RNA-seq from all control and 22q11.2DS hSO samples. Samples clustered by genotype. **(D)** Glycolysis-related genes upregulated in 22q11.2DS: LDHA (Log₂FC = +1.419, p = 0.000078, Wald test, DESeq2), PC (Log₂FC = +1.014, p = 0.054, Wald test, DESeq2), HK2 (Log₂FC = +1.023, p = 0.041, Wald test, DESeq2), PGK1 (Log₂FC = +1.012, p = 0.037, Wald test, DESeq2), PKM (Log₂FC = +0.891, p = 0.0009, Wald test, DESeq2), ENO1 (Log₂FC = +0.875, p = 0.00008, Wald test, DESeq2), SLC2A1 (Log₂FC = +0.868, p = 0.033, Wald test, DESeq2), GAPDH (Log₂FC = +0.817, p = 0.022, Wald test, DESeq2), PFKP (Log₂FC = +0.756, p = 0.0034, Wald test, DESeq2), TPI1 (Log₂FC = +0.674, p = 0.0006, Wald test, DESeq2), PGAM1 (Log₂FC = +0.589, p = 0.041, Wald test, DESeq2), GPI (Log₂FC = +0.584, p = 0.018, Wald test, DESeq2). OXPHOS-related genes in 22q11.2DS upregulated: NDUFAF6 (Log₂FC = +0.805, p = 0.039, Wald test, DESeq2), COX20 (Log₂FC = +0.545, p = 0.029, Wald test, DESeq2), HIGD1A (Log₂FC = +0.395, p = 0.043, Wald test, DESeq2), ATP5PB (Log₂FC = +0.382, p = 0.044, Wald test, DESeq2); downregulated: COX5A (Log₂FC = −0.347, p = 0.012, Wald test, DESeq2), NDUFA6 (Log₂FC = −0.363, p = 0.004, Wald test, DESeq2), ATP5MC1 (Log₂FC = −0.401, p = 0.0018, Wald test, DESeq2), UQCRC2 (Log₂FC = −0.525, p = 0.0031, Wald test, DESeq2), NDUFA1 (Log₂FC = −0.914, p = 0.0025, Wald test, DESeq2), ETFDH (Log₂FC = −0.967, p = 0.012, Wald test, DESeq2). This metabolic shift from oxidative phosphorylation to glycolysis is consistent with the mitochondrial respiration deficits observed in Figure 2. **(E)** Log₂ fold change of genes within the 22q11.2 hemizygously deleted region in 22q11.2DS versus control hSOs: PRODH, TANGO2, MRPL40, ZDHHC8, TXNRD2, SLC25A1, COMT, DGCR8. Most genes show the expected ∼0.5-fold reduction consistent with hemizygous deletion. Data are represented as mean ± SEM.

**Figure S3.**
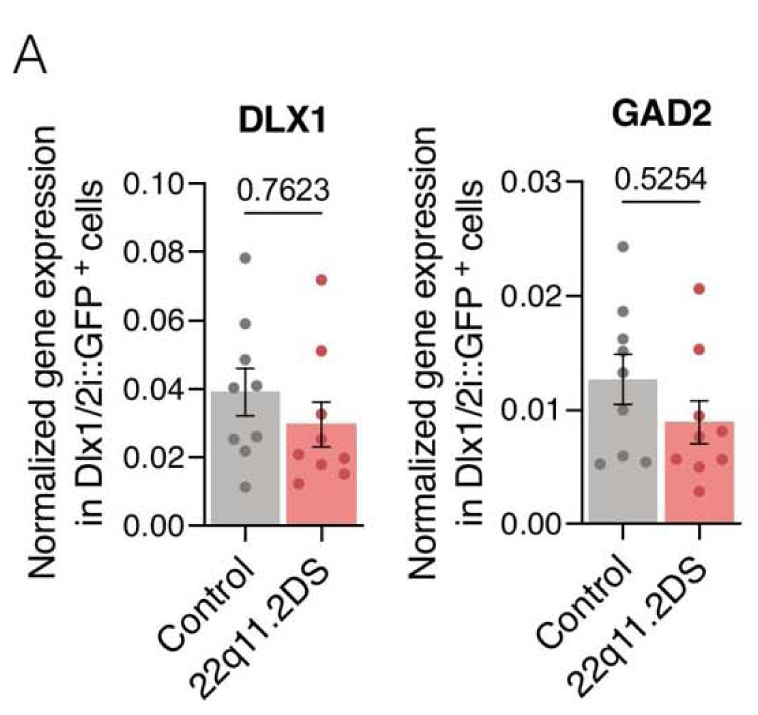
Normalized Gene Expression of Inhibitory Neuron Markers in Sorted Dlx1/2i::GFP Positive Cells, Related to. Figure 2 **Interneuron identity markers are comparably expressed in control and 22q11.2DS Dlx1/2i::GFP+ cells, confirming that migration deficits are not attributable to altered cell fate. (A)** qRT-PCR of *DLX1* (p = 0.7623; Welch’s t-test) and *GAD2* (p = 0.5254; Welch’s t-test) in Dlx1/2i::GFP+ cells. Each dot represents one biological replicate: *n* = 4 iPSC lines per genotype. Data are represented as mean ± SEM.

**Figure S4.**
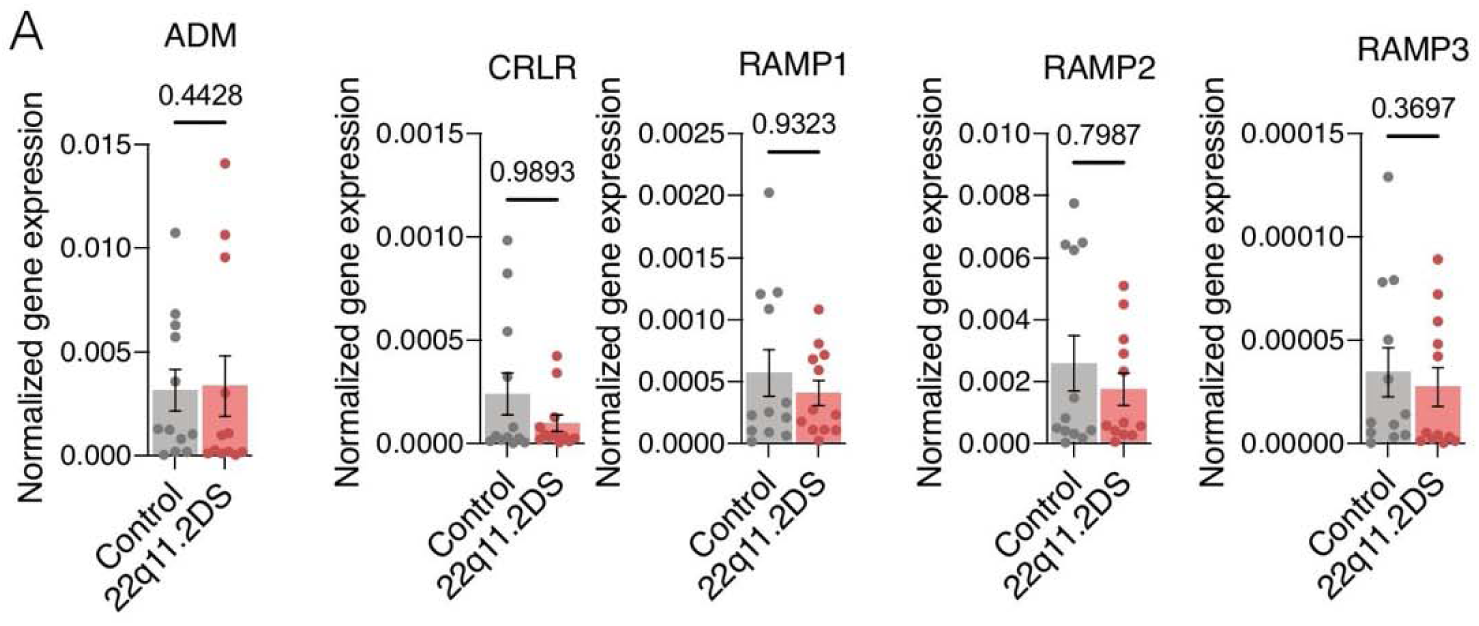
ADM Receptor Expression in 22q11.2DS hSOs, Related to Figure 3 Expression of *ADM* and its receptor components is comparable between genotypes, confirming intact signaling machinery in 22q11.2DS hSOs. (A) qRT-PCR of *ADM* (p = 0.4428; Welch’s t-test), *CRLR* (p = 0.9893; Welch’s t-test), *RAMP1* (p = 0.9323; Welch’s t-test), *RAMP2* (p = 0.7987; Welch’s t-test), and *RAMP3* (p = 0.3697; Welch’s t-test). Each dot represents a biological replicate; *n* = 3 per genotype. Data are represented as mean ± SEM.

**Figure S5.**
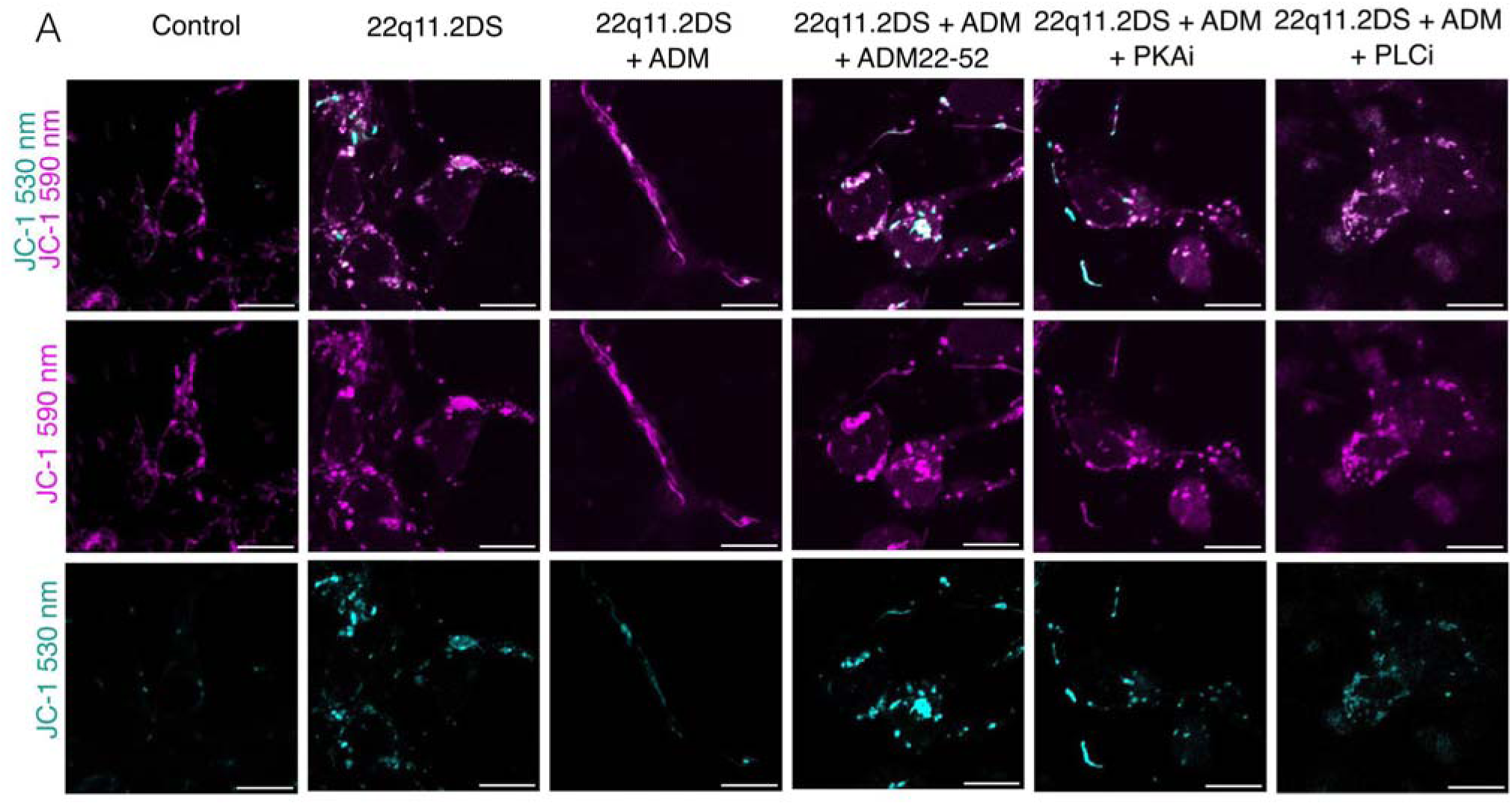
JC-1 Channel Separation across Pharmacological Conditions, Related to. Figure 6 **Individual JC-1 fluorescence channels confirm that ADM restores mitochondrial membrane potential in 22q11.2DS hSOs and that this effect is blocked by receptor antagonism (ADM22-52), PKA inhibition, and PLC inhibition. (A)** Full JC-1 imaging panels across six conditions: control, 22q11.2DS, 22q11.2DS + ADM, 22q11.2DS + ADM + ADM22-52, 22q11.2DS + ADM + PKAi, and 22q11.2DS + ADM + PLCi. Row 1: JC-1 530 nm + 590 nm merged. Row 2: JC-1 590 nm only (magenta, J-aggregates indicating high MMP). Row 3: JC-1 530 nm only (cyan, monomers indicating low MMP). Scale bar, 20 µm.

