## Supplementary material for "Adrenomedullin Restores Mitochondrial Bioenergetics and Rescues Interneuron Phenotypes in Human Models of 22q11.2 Deletion Syndrome": Table S2

**SUPPLEMENTARY TABLE 2**

Primers used for RT-PCR experiments

| **Gene** | **Forward Primer** | **Reverse Primer** |
| --- | --- | --- |
| GAPDH | GGAGCGAGATCCCTCCAAAAT | GGCTGTTGTCATACTTCTCATGG |
| GAD2 | GCCAACTCTGTGACGTGGAATC | GCTGAAAGAGGTAGGAGGCATG |
| LHX6 | CGCATCCACTACGACACCATGA | GCTTGGGTTGACTGTCCTGTTC |
| CALB | AGGGAATCAAAATGTGTGGGAAA | TCCTTCAGTAAAGCATCCAGTTC |
| SST | GCTGCTGTCTGAACCCAAC | CGTTCTCGGGGTGCCATAG |
| PVALB | GGACAAAAGTGGCTTCATCGAG | TCGTCAACCCCAATTTTGCC |
| FOXG1 | AACCTGTGTTGCGCAAATGC | AAACACGGGCATATGACCAC |
| EMX1 | CGCAGGTGAAGGTGTGGTT | TCCAGCTTCTGCCGTTTGT |
| DLX1 | ATGCACTGTTTACACTCGGC | GACTGCACCGAACTGATGTAG |
| ADM | GAACTGCGGATGTCCAG | GTCCTTGTCCTTATCTGTG |
| CRLR | TGGATGGCTCTGCTGGAACGAT | CCAGTTTCCATCTTGGTCACAGA |
| RAMP1 | CTCACCCAGTTCCAGGTAGACA | CAGGAAGAACCTGTCCACCTCT |
| RAMP2 | AGCTGTCCAATTTTGCTGGAATCA | CAAAGTGCTCCAGGCAATCTCG |
| RAMP3 | ACGTCTGGAAGTGGTGCAACCT | CCAGTAGCAGCCCACGACATTG |
