## Supplementary material for "Adrenomedullin Restores Mitochondrial Bioenergetics and Rescues Interneuron Phenotypes in Human Models of 22q11.2 Deletion Syndrome": Table S3

| **Table S3. Mitochondrial Process Dysregulation in 22q11.2DS vs Control hSOs** | | | | | |
| --- | --- | --- | --- | --- | --- |
| **Mitochondrial Process** | **Gene** | **Direction** | **log2FC** | **P-value** | **FDR** |
| Amino Acid Metabolism | **ETFDH** | **Down** | -0.967 | 7.82E-04 | 2.45E-01 |
| Amino Acid Metabolism | **COMT** | **Up** | +1.446 | 1.04E-03 | 2.45E-01 |
| Amino Acid Metabolism | **HSD17B10** | **Up** | +0.755 | 1.47E-02 | 4.13E-01 |
| Amino Acid Metabolism | **GCSH** | **Up** | +0.679 | 3.60E-03 | 2.79E-01 |
| Amino Acid Metabolism | **ECHS1** | **Up** | +0.567 | 2.39E-02 | 4.63E-01 |
| Apoptosis | **BBC3** | **Down** | -0.682 | 2.90E-02 | 4.86E-01 |
| Apoptosis | **BNIP3** | **Up** | +1.201 | 7.15E-05 | 1.31E-01 |
| Apoptosis | **IFI27** | **Up** | +1.019 | 4.32E-03 | 2.98E-01 |
| Apoptosis | **RTL10** | **Up** | +0.954 | 3.12E-03 | 2.73E-01 |
| Calcium Signaling & Transport | **RHOT2** | **Down** | -0.611 | 1.61E-02 | 4.18E-01 |
| Calcium Signaling & Transport | **EFHD1** | **Down** | -1.179 | 4.85E-03 | 3.00E-01 |
| Cardiolipin Biosynthesis | **PTPMT1** | **Up** | +0.998 | 7.32E-03 | 3.28E-01 |
| Fatty Acid Biosynthesis | **HTD2** | **Down** | -0.674 | 3.81E-02 | 5.16E-01 |
| Fatty Acid Metabolism | **ETFDH** | **Down** | -0.967 | 7.82E-04 | 2.45E-01 |
| Fatty Acid Metabolism | **HSD17B10** | **Up** | +0.755 | 1.47E-02 | 4.13E-01 |
| Fatty Acid Metabolism | **ECHS1** | **Up** | +0.567 | 2.39E-02 | 4.63E-01 |
| Fe-S Cluster Biosynthesis | **FDXR** | **Up** | +0.842 | 2.64E-02 | 4.69E-01 |
| Glycolysis | **LDHA** | **Up** | +1.419 | 7.80E-05 | 1.31E-01 |
| Glycolysis | **HK2** | **Up** | +1.023 | 4.09E-02 | 5.23E-01 |
| Glycolysis | **PGK1** | **Up** | +1.012 | 8.00E-05 | 1.31E-01 |
| Glycolysis | **PKM** | **Up** | +0.891 | 9.02E-04 | 2.45E-01 |
| Glycolysis | **ENO1** | **Up** | +0.875 | 2.03E-03 | 2.70E-01 |
| Glycolysis | **SLC2A1** | **Up** | +0.868 | 6.87E-03 | 3.28E-01 |
| Glycolysis | **GAPDH** | **Up** | +0.817 | 2.40E-03 | 2.70E-01 |
| Glycolysis | **PFKP** | **Up** | +0.756 | 8.17E-03 | 3.34E-01 |
| Glycolysis | **TPI1** | **Up** | +0.674 | 7.22E-03 | 3.28E-01 |
| Glycolysis | **PGAM1** | **Up** | +0.589 | 1.95E-02 | 4.39E-01 |
| Glycolysis | **GPI** | **Up** | +0.584 | 5.52E-03 | 3.14E-01 |
| Import & Sorting | **UQCRC2** | **Down** | -0.525 | 2.15E-02 | 4.54E-01 |
| Import & Sorting | **AGK** | **Down** | -0.755 | 1.99E-02 | 4.39E-01 |
| Import & Sorting | **TIMM10B** | **Down** | -0.791 | 1.69E-02 | 4.21E-01 |
| Import & Sorting | **TIMM10** | **Up** | +0.676 | 1.29E-02 | 3.90E-01 |
| Import & Sorting | **TIMM8B** | **Up** | +0.661 | 7.28E-03 | 3.28E-01 |
| Import & Sorting | **TIMM23** | **Up** | +0.502 | 3.23E-02 | 4.97E-01 |
| Mitochondrial Carrier | **SLC25A46** | **Down** | -0.593 | 2.85E-02 | 4.85E-01 |
| Mitochondrial Carrier | **SLC25A42** | **Down** | -0.786 | 4.77E-02 | 5.46E-01 |
| Mitochondrial Carrier | **SLC25A1** | **Up** | +0.734 | 4.93E-03 | 3.01E-01 |
| Mitochondrial Dynamics | **PARL** | **Down** | -0.568 | 1.96E-02 | 4.39E-01 |
| Mitochondrial Dynamics | **SLC25A46** | **Down** | -0.593 | 2.85E-02 | 4.85E-01 |
| Mitochondrial Dynamics | **RHOT2** | **Down** | -0.611 | 1.61E-02 | 4.18E-01 |
| Mitochondrial Dynamics | **BBC3** | **Down** | -0.682 | 2.90E-02 | 4.86E-01 |
| Mitochondrial Dynamics | **MTFR1** | **Down** | -0.843 | 4.96E-02 | 5.48E-01 |
| Mitochondrial Dynamics | **SNAP29** | **Up** | +1.257 | 1.38E-04 | 1.36E-01 |
| Mitochondrial Dynamics | **BNIP3** | **Up** | +1.201 | 7.15E-05 | 1.31E-01 |
| Mitochondrial Dynamics | **IFI27** | **Up** | +1.019 | 4.32E-03 | 2.98E-01 |
| Mitochondrial Dynamics | **RTL10** | **Up** | +0.954 | 3.12E-03 | 2.73E-01 |
| Mitochondrial Signaling | **RHOT2** | **Down** | -0.611 | 1.61E-02 | 4.18E-01 |
| Mitochondrial Signaling | **EFHD1** | **Down** | -1.179 | 4.85E-03 | 3.00E-01 |
| Mitochondrial Signaling | **IFI27** | **Up** | +1.019 | 4.32E-03 | 2.98E-01 |
| Oxidative Phosphorylation | **UQCRC2** | **Down** | -0.525 | 2.15E-02 | 4.54E-01 |
| Oxidative Phosphorylation | **NDUFA1** | **Down** | -0.914 | 2.97E-02 | 4.91E-01 |
| Oxidative Phosphorylation | **NDUFAF8** | **Up** | +0.836 | 9.52E-03 | 3.49E-01 |
| Oxidative Phosphorylation | **NDUFAF6** | **Up** | +0.805 | 1.88E-02 | 4.33E-01 |
| Oxidative Phosphorylation | **COX20** | **Up** | +0.545 | 3.18E-02 | 4.97E-01 |
| Pyruvate Metabolism | **PDK1** | **Up** | +0.836 | 1.46E-02 | 4.13E-01 |
| ROS Defense | **GLRX2** | **Down** | -1.493 | 3.01E-03 | 2.73E-01 |
| ROS Defense | **MGST1** | **Up** | +1.284 | 1.89E-03 | 2.66E-01 |
| ROS Defense | **TXNRD2** | **Up** | +1.017 | 4.14E-02 | 5.24E-01 |
| Replication & Transcription | **RECQL4** | **Up** | +0.904 | 3.64E-02 | 5.08E-01 |
| Translation | **MRPL2** | **Down** | -0.593 | 3.97E-02 | 5.23E-01 |
| Translation | **MRPL10** | **Down** | -0.674 | 4.31E-02 | 5.33E-01 |
| Translation | **MRPL40** | **Up** | +1.018 | 4.21E-04 | 2.09E-01 |
