## Supplementary material for "Adrenomedullin Restores Mitochondrial Bioenergetics and Rescues Interneuron Phenotypes in Human Models of 22q11.2 Deletion Syndrome": Key Resource Table

| **REAGENT or RESOURCE** | **SOURCE** | **IDENTIFIER** |
| --- | --- | --- |
| **Antibodies** | | |
| β-actin antibody (anti-mouse, Clone 13E5) | Cell Signaling Technology | Cat#4970S |
| Anti-SOX2 (goat anti-human) | R&D Systems | Cat# AF2018 |
| Anti-DRP1 (Total) (Rabbit monoclonal) | Cell Signaling Technology | Cat# 85705 |
| Anti-Phospho-DRP1 (Ser637) (Rabbit polyclonal) | Cell Signaling Technology | Cat# 4867 |
| Anti-Phospho-DRP1 (Ser616) (Rabbit polyclonal) | Cell Signaling Technology | Cat# 3455 |
| Anti-TUJ1 (Mouse monoclonal) | Biolegend | Cat# 801202 |
| Anti-mouse IgG HRP-conjugated secondary antibody | Cell Signaling Technology | Cat#7076 |
| Anti-rabbit IgG HRP-conjugated secondary antibody | Cell Signaling Technology | Cat#7074 |
| Donkey anti-Goat IgG (H+L) Cross-Adsorbed Secondary Antibody, Alexa Fluor™ 488 | Invitrogen | Cat# A-11055 |
| Donkey anti-Mouse IgG (H+L) Highly Cross-Adsorbed Secondary Antibody, Alexa Fluor™ 647 | Invitrogen | Cat# A-31571 |
| Zombie Violet™ Fixable Viability Kit | Biolegend | Cat# 423113 |
| **Bacterial and virus strains** | | |
| Lentivirus: Lenti-Dlxi1/2b::eGFP | Gift from J.L. Rubenstein lab | N/A |
| **Biological samples** | | |
| Human cortical tissue (20 PCW) | Stanford University (IRB-approved protocol) | N/A |
| **Chemicals, peptides, and recombinant proteins** | | |
| Human Adrenomedullin (ADM, 1-52) peptide | Anaspec | Cat#AS-60447 |
| ADM22-52 (AM receptor antagonist) | Cayman Chemicals | Cat#24892 |
| KT5720 (PKA inhibitor) | Tocris Bioscience | Cat#1288 |
| U-73122 (PLC inhibitor) | Tocris Bioscience | Cat#1268 |
| Oligomycin A | Sigma-Aldrich | Cat#75351 |
| FCCP (carbonyl cyanide 4-(trifluoromethoxy)phenylhydrazone) | Sigma-Aldrich | Cat#C2920 |
| Rotenone | Cayman Chemicals | Cat#13995 |
| Dorsomorphin (2.5 µM) | Sigma-Aldrich | Cat#P5499 |
| SB-431542 (10 µM) | Tocris | Cat#1614 |
| XAV-939 | Tocris | Cat#3748 |
| SAG (Smoothened Agonist, 100 nM) | Thermo Fisher Scientific | Cat#566660 |
| CEPT cocktail | Selleck Chemicals | Cat#S1049 |
| Calbryte™ 520 AM (calcium indicator) | AAT BioQuest | Cat#20650 |
| PKmito DEEP RED (mitochondrial probe) | Spirochrome | Cat#SC055 |
| JC-1 Dye (mitochondrial membrane potential probe) | Thermo Fisher Scientific | Cat#T3168 |
| SPY555-FastAct™ (live-cell F-actin probe) | Spirochrome | Cat#SC205 |
| EGF, recombinant human (20 ng/mL) | R&D Systems | Cat#236-EG |
| FGF2, recombinant human (20 ng/mL) | R&D Systems | Cat#233-FB |
| BDNF, recombinant human (20 ng/mL) | Peprotech | Cat#450-02 |
| NT3, recombinant human (20 ng/mL) | Peprotech | Cat#450-03 |
| Recombinant Human Vitronectin (VTN-N) | Thermo Fisher Scientific | Cat#A14700 |
| Essential 8 Medium | Thermo Fisher Scientific | Cat#A1517001 |
| Essential 6 Medium | Thermo Fisher Scientific | Cat#A1516401 |
| Neurobasal A | Thermo Fisher Scientific | Cat#10888022 |
| B-27 Supplement, without Vitamin A (50X) | Thermo Fisher Scientific | Cat#12587010 |
| GlutaMax (200 mM, 100X) | Thermo Fisher Scientific | Cat#35050061 |
| Penicillin-Streptomycin (100X) | Thermo Fisher Scientific | Cat#15140122 |
| N-2 Supplement (100X) | Gibco | Cat#17502048 |
| Fetal Bovine Serum (FBS), certified | Gibco | Cat#16000044 |
| Basal Medium Eagle (BME) | Gibco | Cat#21010046 |
| Hanks' Balanced Salt Solution (HBSS) | Gibco | Cat#14025092 |
| D-(+)-Glucose, BioReagent, ≥99.5% | Sigma-Aldrich | Cat#G7021 |
| DPBS (Dulbecco's Phosphate Buffered Saline) | Thermo Fisher Scientific | Cat#14190144 |
| Tween 20 | Sigma-Aldrich | Cat#P1379 |
| BSA (bovine serum albumin) | Gendepot | Cat#A0100-005 |
| Skim milk powder | BD | Cat#232100 |
| Tris-Buffered Saline (TBS) | Boston BioProducts | Cat#BM-301X |
| SuperSignal™ West Femto Maximum Sensitivity Substrate | Thermo Fisher Scientific | Cat#34095 |
| UltraPure Low Melting Point Agarose | Invitrogen | Cat#16520-100 |
| RIPA buffer with protease and phosphatase inhibitor cocktail | Santacruz | Cat#sc-24948A |
| Bolt™ LDS Sample Buffer | Invitrogen | Cat#B0007 |
| Bolt™ 4–12% Bis-Tris Protein Gels | Invitrogen | Cat#NW04120BOX |
| PEI Max (transfection reagent) | Polysciences | Cat#24765 |
| **Critical commercial assays** | | |
| RNeasy Mini Kit with RNase-Free DNase Set | Qiagen | Cat#74136 |
| SuperScript IV First-Strand Synthesis System | Thermo Fisher Scientific | Cat# 18091050 |
| PowerUp™ SYBR™ Green Master Mix | Life Technologies | Cat#A25742 |
| Pierce™ BCA Protein Assay Kit | Thermo Fisher Scientific | Cat#23225 |
| iBlot™ 2 Transfer Stacks (PVDF membrane) | Invitrogen | Cat#IB24002 |
| **Experimental models: Cell lines** | | |
| Human: hiPSC control lines (for hCO, hSO, hFA generation) | Pasca lab, Stanford University | N/A |
| Human: hiPSC 22q11.2DSl lines (for hCO, hSO, hFA generation) | Pasca lab, Stanford University | N/A |
| Human: HEK293T | ATCC | Cat# CRL-3216, RRID:CVCL_0063 |
| **Oligonucleotides** | | |
| Primers for qRT-PCR, see Table S2 | This paper | N/A |
| **Software and algorithms** | | |
| NovoExpress (flow cytometry software) | Agilent | v1.5.6; https://www.agilent.com |
| FlowJo (flow cytometry analysis) | BD Biosciences | v10.8.1; https://www.flowjo.com |
| BioRad CFX Maestro (qPCR data processing) | BioRad | https://www.bio-rad.com |
| ImageJ (western blot band quantification) | NIH/NIMH | v1.53t; https://imagej.nih.gov/ij/ |
| GraphPad Prism 10 | GraphPad Software | RRID:SCR_002798 |
| MaxLive Software | Maxwell Biosystems | N/A |
| XF Software Agilent Seahorse Analytics | Agilent | https://seahorseanalytics.agilent.com/ |
| **Other** | | |
| AggreWell 800 (organoid formation) | STEMCELL Technologies | Cat#34815 |
| Ultra-low-attachment dishes | Corning | Cat#430293 |
| 24-well ultra-low-attachment plate | Corning | Cat#3474 |
| 96-well glass-bottom plate (live imaging) | Cellvis | Cat#P96-0N |
| Cell culture membrane inserts (0.4 µm pore, 23.1 mm diameter) | Falcon | Cat#353090 |
| 40 µm cell strainer | Biologix Research Company | Cat#15-1040 |
| Zeiss LSM 980 confocal microscope with motorized stage | Zeiss | N/A |
| Okolab Bold Line cage incubator | Okolab | N/A |
| Leica VT1200 vibratome | Leica | N/A |
| NovoCyte Penteon flow cytometer (Stanford FACS Facility) | Agilent | N/A |
| CFX384 real-time qPCR system | BioRad | N/A |
| iBright 1500 imaging system | Thermo Fisher Scientific | Cat#A44114 |
| iBlot 2 dry blotting system | Thermo Fisher Scientific | Cat#IB21001 |
| Seahorse xFe96 Analyzer | Agilent | N/A |
| **Other** | | |
| Raw and processed scRNA-seq data: forebrain assembloids (day 60+), migrating inhibitory interneurons | Walsh et al., 2026 | GEO: GSE250482 |
| **Software and Algorithms** | | |
| CellAtria (agentic AI data ingestion and orchestration framework) | Nouri et al., 2026 | v1.0.0; https://github.com/AstraZeneca/cellatria |
| CellExpress (automated scRNA-seq processing pipeline) | Nouri et al., 2026 | v1.0.0; https://github.com/AstraZeneca/cellatria |
| Scanpy | Wolf et al., 2018 | ≥1.9; https://scanpy.readthedocs.io |
| Scrublet | Wolock et al., 2019 | https://github.com/swolock/scrublet |
| Harmony | Korsunsky et al., 2019 | https://github.com/immunogenomics/harmony |
| SCimilarity | Heimberg et al., 2023 | v1.1; https://github.com/Genentech/scimilarity |
| CellTypist | Dominguez Conde et al., 2022 | v1.6.3; https://www.celltypist.org |
| UMAP | McInnes et al., 2018 | https://github.com/lmcinnes/umap |
| Leiden algorithm | Traag et al., 2019 | https://leidenalg.readthedocs.io |

**Declaration of generative AI and AI-assisted technologies in the manuscript preparation process**

During the preparation of this work, the authors used Grammarly to improve the text's readability. After using this tool, the authors reviewed and edited the content as needed and take full responsibility for the content of the published article.
